# FUS-ALS mutants alter FMRP phase separation equilibrium and impair protein translation

**DOI:** 10.1101/2020.09.14.296038

**Authors:** N. Birsa, A.M. Ule, M.G. Garone, B. Tsang, F. Mattedi, P.A. Chong, J. Humphrey, S. Jarvis, M. Pisiren, O.G. Wilkins, M. Nosella, A. Devoy, C. Bodo, R. Fernandez de la Fuente, E.M.C. Fisher, A. Rosa, G. Viero, J.D. Forman-Kay, G. Schiavo, P. Fratta

**Author notes:** corresponding authors: Pietro Fratta; Nicol Birsa. these authors have contributed equally to this manuscript (N. Birsa, A.M. Ule). these authors have contributed equally to this manuscript (M.G. Garone, B. Tsang, F. Mattedi).

## Abstract

Mutations in the RNA binding protein (RBP) FUS cause amyotrophic lateral sclerosis (ALS) and result in its nuclear depletion and cytoplasmic mislocalisation, with cytoplasmic gain of function thought to be crucial in pathogenesis. Here, we show that expression of mutant FUS at physiological levels drives translation inhibition in both mouse and human motor neurons. Rather than acting directly on the translation machinery, we find that mutant FUS forms cytoplasmic condensates that promote the phase separation of FMRP, another RBP associated with neurodegeneration and robustly involved in translation regulation. FUS and FMRP co-partition and repress translation *in vitro*. In our *in vivo* model, FMRP RNA targets are depleted from ribosomes. Our results identify a novel paradigm by which FUS mutations favour the condensed state of other RBPs, impacting on crucial biological functions, such as protein translation.

## Introduction

Amyotrophic lateral sclerosis (ALS) is a devastating neurodegenerative disorder caused by the progressive loss of motor neurons (MNs), leading to muscle weakness and ultimately death by respiratory failure. Mutations in genes encoding several RNA binding proteins (RBPs), including TAR DNA Binding Protein 43 (TDP-43) and FUsed in Sarcoma (FUS), are causative of ALS (Mejzini et al., 2019; Taylor et al., 2016), suggesting that perturbations in RNA metabolism may play a pivotal role in disease pathogenesis. Mutations in FUS account for approximately 5% of familial ALS, including cases with very early onset and aggressive disease course (Taylor et al., 2016). In physiological conditions, FUS is predominantly localised in the nucleus where it is involved in transcription, mRNA processing and microRNA biogenesis (Ederle and Dormann, 2017). Low levels of the protein are also present in the cytoplasm, where FUS has a role in mRNA stability, transport and the stress response (Birsa et al., 2020).

The majority of FUS ALS-causative mutations disrupt the C-terminal nuclear localisation signal (NLS) leading to a depletion from the nucleus and a cytoplasmic mislocalisation of the protein (Dormann et al., 2010); further, cytoplasmic FUS-positive aggregates are present in patient *post mortem* tissue (Vance et al., 2009). An outstanding question, critical to mechanistically address ALS, is whether it is a nuclear loss of function, a cytoplasmic gain of function or a combination of the two that drives mutant FUS-mediated toxicity.

We have recently shown that, unlike TDP-43, where ALS-associated mutations cause a nuclear gain of function (Fratta et al., 2018), mutations in FUS result in changes that are in line with a nuclear loss of function (Humphrey et al., 2020). However, in contrast to models expressing FUS mutants at physiological levels (Devoy et al., 2017; Scekic-Zahirovic et al., 2017), FUS knock-out animals do not show overt MN loss and functional motor impairment (Kino et al., 2015) supporting a toxic cytoplasmic gain of function to have a role in disease pathogenesis.

The role of FUS in the cytoplasm and the functional consequences of ALS-linked mutations are poorly understood. In addition to a role in stress response, it was recently observed that, in transgenic and overexpression models, mutant FUS was associated with an impairment in protein translation (Kamelgarn et al., 2018; López-Erauskin et al., 2018; Murakami et al., 2015). Further, an increasing number of studies suggests that FUS liquid-liquid phase separation (LLPS) properties play a crucial role in its cytoplasmic gain of function. Cytoplasmic condensates, such as stress granules and RNA granules, are generated by phase separation and have various functional roles, generally bringing together specific sets of proteins, RNAs and other factors. In physiological conditions, the propensity of FUS to undergo LLPS in the cytoplasm is limited by its low concentration levels. In contrast, the increased cytoplasmic localisation of FUS mutants favours this transition. Moreover, a decreased interaction with the nuclear import factor and chaperone TNPO1, post-translational modifications and intrinsic properties given by ALS mutations are all factors contributing to the formation of cytoplasmic FUS condensates (Hofweber et al., 2018; Murakami et al., 2015; Qamar et al., 2018), the biology and composition of which are, to date, poorly understood. Importantly, potential disease mechanisms have been identified in various overexpression models (Murakami et al., 2015; Patel et al., 2015), however, it remains unclear how well these model the physiological disease setting.

Here we use mouse and human models with endogenous expression of FUS-ALS mutations and show that mutant FUS induces a decrease in global protein translation. We find that this is not caused by direct interaction with the translational machinery; rather mutant FUS forms cytoplasmic condensates containing the RBP Fragile X Mental Retardation Protein (FMRP), resulting in decreased translation of FMRP mRNA targets. Our findings highlight how, in addition to a previously described effect at the transcriptional level, mutant FUS can post-transcriptionally alter RBP dynamics and function.

## Results

### 1. Cytoplasmic FUS represses translation

Two reports have recently shown that mutant FUS overexpression can impair protein synthesis in neurons (López-Erauskin et al., 2018; Murakami et al., 2015), but whether this also occurs with physiological FUS expression is not currently known. In order to investigate this, we used the Δ14 FUS knock-in mouse model (Devoy et al., 2017), in which a mutation causing aggressive and early-onset ALS (DeJesus-Hernandez et al., 2010), leads to skipping of exon 14 and a frameshift in exon 15. This frameshift results in the loss of the entire NLS and the generation of a unique C-terminal region (SFig. 1A), which has allowed us to generate mutant-specific antibodies (Devoy et al., 2017). Similar to ΔNLS FUS homozygous mice (Scekic-Zahirovic et al., 2016), Δ14 FUS homozygotes die at birth (Devoy et al., 2017). Therefore, to investigate the function of Δ14 FUS in MNs, as well as its dosage-dependent regulation, we used primary embryonic cultures. While wildtype FUS has a predominantly nuclear localisation in both *Fus*^+/+^ and *Fus*^Δ14/+^ MNs, Δ14 FUS is enriched in the cytoplasm of *Fus*^Δ14/+^ and *Fus*^Δ14/Δ14^ neurons, it is homogeneously distributed throughout the cell body and neurites, it can be detected in a punctate, condensate-like form and no large inclusions have been observed (Fig. 1A,B; SFig. 1B-D). Of note, Δ14 FUS expression in *Fus*^Δ14/+^ MNs did not induce the mislocalisation of wildtype FUS (Fig. 1A; SFig. 1B,C).

**Fig. 1.**
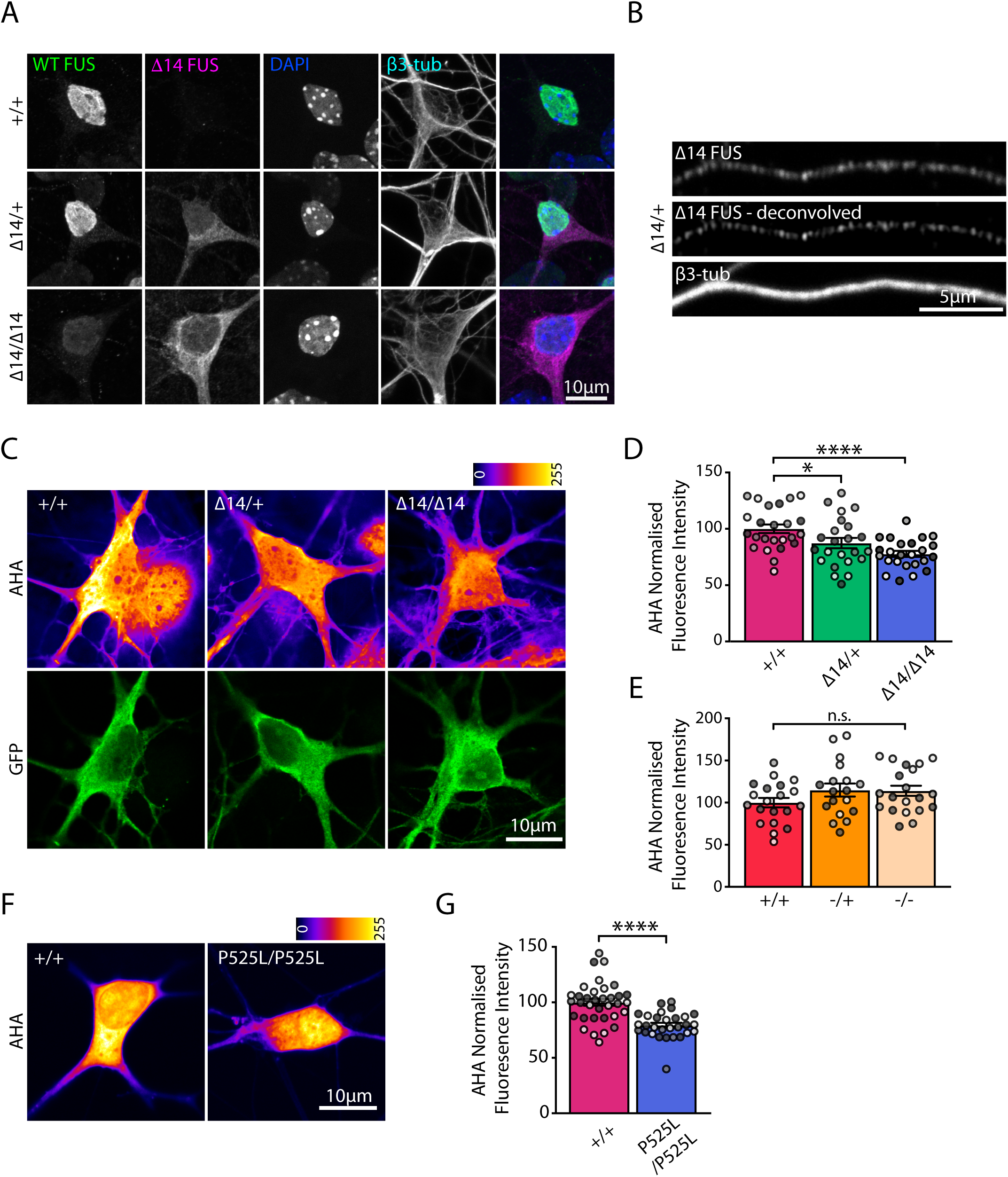
Protein translation is reduced in mutant FUS expressing MNs. **(A)** Representative images of primary motor neurons (DIV5). Wild type FUS (green), detected with a C-term antibody, is primarily localised in the nucleus in *Fus*^+/+^ and *Fus*^Δ14/+^ neurons. Δ14 FUS (magenta) is enriched in the cytoplasm of *Fus*^Δ14/+^ and *Fus*^Δ14/Δ14^ neurons. Nuclei are labelled with DAPI (blue) and β3-tubulin is used as a neuronal marker. Low intensity wild type FUS-positive nuclear staining in *Fus*^Δ14/Δ14^ neurons is due to antibody cross reactivity with another FET protein, likely EWSR1. **(B)** Δ14 FUS distribution in a *Fus*^Δ14/+^ MN axon. Δ14 FUS signal detected by confocal microscopy (top panel) and the deconvoluted signal (middle panel). Neurons were grown in microfluidic devices and β3-tubulin is used to identify axons. **(C)** Representative images of primary *Fus*^+/+^, *Fus*^Δ14/+^ and *Fus*^Δ14/Δ14^ MNs metabolically labelled using the methionine analog L-azidohomoalanine (AHA, 2 mM, 30 min) and click chemistry. AHA labelling is visualised using the LUT fire (top panels), motor neurons are identified by GFP expression under the HB9 promoter (bottom panels). **(D)** Quantification of the AHA labelling as shown in (C). Mean fluorescence intensity values are normalised to *Fus*^+/+^ (n=4, MNs=27-28, one way ANOVA followed by Dunnett’s multiple comparisons test, *p<0.05, ****p<0.0001). **(E)** Quantification of translation assays carried out in *Fus*^+/+^, *Fus*^+/-^ and *Fus*^-/-^ motor neurons. Mean fluorescence intensity values are normalised to *Fus*^+/+^ (n=3, MNs=18-20). **(F)** Representative images of isogenic control (*FUS^+/+^*) and *FUS*^P525L/P525L^ iPSC-derived MNs metabolically labelled with AHA (2 mM, 30 min). **(E)** Quantification of the effect in (F). Fluorescence values are normalised to *FUS^+/+^* MNs (n=3, MNs=29-34; Student’s t-test ****p<0.0001). Independent experiments are visualised in different shades of grey in the graphs.

To analyse *de-novo* protein synthesis, we used the methionine analog L-azidohomoalanine (AHA, 2 mM, 30 min incubation) in combination with click chemistry, and found that compared to wildtype controls, AHA labelling was reduced by 13% in *Fus*^Δ14/+^ and by 20% in *Fus*^Δ14/Δ14^ primary MNs (Fig. 1C,D; average AHA intensity in *Fus*^+/+^ MNs 100±3.8, *Fus*^Δ14/+^ MNs 87.4±4.5 *Fus*^Δ14/Δ14^ MNs 77.7±2.7; *p<0.05, ****p<0.0001). As a positive control we showed that pre-treatment with the translation inhibitor anisomycin (40 μM, 20 min) reduced the AHA signal by 80%, indicating the specificity of the labelling (SFig. 2B,C). In contrast, in FUS knock-out MNs no effect on *de-novo* protein synthesis was detected (Fig. 1E; SFig. 2A; average AHA intensity in *Fus*^+/+^ MNs 100±5.5, *Fus*^+/-^ MNs 115±7.7, *Fus*^-/-^ MNs 114±6.0), indicating that inhibition of translation is due to a gain of function of mutant FUS.

To test whether this deficit was conserved in human models of FUS-ALS, we performed the same assay in iPSC-derived MNs carrying the common ALS-associated NLS mutation P525L and compared them to isogenic controls. With a remarkable similarity to the primary mouse MNs, AHA labelling was decreased by 20% in mutant FUS-expressing human MNs (Fig. 1F,G, isogenic control 100±3.1, *FUS*^P525L/P525L^ 79.4±2.2; ****p<0.0001), underlining that translation inhibition stems from FUS cytoplasmic mislocalisation and is not mutation-specific.

### 2. Mutant FUS does not alter translation by association with polysomes

Despite its low cytoplasmic levels, wildtype FUS binds proteins of both the small and large ribosomal subunits (Simsek et al., 2017), suggesting that it may indeed interact with assembled ribosomes. Given the increase in cytoplasmic levels of mutant FUS, we asked whether association with ribosomes could account for the observed changes in translation. To investigate this, we performed polysome co-sedimentation assays, where separation of the heavier polysomal fractions from monosomes (80S), the individual ribosomal subunits (60S and 40S) and the lighter free cytosolic complexes (SFig. 3Ai) allows the analysis of the association of specific proteins with the translation components. As expected, the majority of wildtype FUS co-sedimented with free cytosolic complexes, but significant levels also co-sedimented with polysomes in both *Fus*^+/+^ and *Fus*^Δ14/+^ samples (SFig. 3Aii). Instead, despite its largely cytoplasmic localisation, Δ14 mutant FUS did not co-sediment with polysomal fractions (fractions 8-11; SFig. 3Aii) and was only present in the lighter part of the gradient, containing sub-polysomal components, in *Fus*^Δ14/+^ and *Fus*^Δ14/Δ14^ samples (fractions 1-6; SFig. 3Aii).

FUS interacts with several other RBPs, among which FMRP and SMN are the most well-characterised and are strongly linked to pathologies of the nervous system (Blokhuis et al., 2016; Groen et al., 2013; Sun et al., 2015; Yamazaki et al., 2012). As both RBPs are also known to associate with the translational machinery (Bernabò et al., 2017; Darnell et al., 2011), we examined their co-sedimentation profile in our Δ14 FUS model. While both FMRP and SMN interact with Δ14 FUS (SFig. 4A), when analysing their co-sedimentation, we found that the distribution of SMN was unaltered by the expression of mutant FUS, whereas FMRP, similarly to mutant FUS itself, was depleted from the polysomal fractions in both *Fus*^Δ14/+^ and *Fus*^Δ14/Δ14^ samples (SFig. 3A). We confirmed the weaker association of FMRP with ribosomes in Δ14 FUS MNs using proximity ligation assays (PLA) between FMRP and the ribosomal protein RPL26 (SFig. 3C,D; normalised FMRP-RPL26 PLA puncta in *Fus*^+/+^=1.0±0.14, *Fus*^Δ14/+^=0.18±0.04, *Fus*^Δ14/Δ14^=0.37±0.1; *p<0.05, ****p<0.0001). Importantly, we found FUS depletion (*Fus*^-/-^) to have a similar effect on FMRP-ribosome association both in co-sedimentation assays (SFig. 3B), and in FMRP-RPL26 PLA (SFig. 3E; normalised FMRP-RPL26 PLA puncta in *Fus*^+/+^=1.0±0.12, *Fus*^+/-^ =0.9±0.21, *Fus*^-/-^=0.44±0.08; **p<0.001).

In summary, these results show that mutations in FUS impair the association of the protein with polysomes and that wildtype FUS has a role in influencing the association of FMRP with the translation machinery. Given that the global level of protein synthesis is not affected in FUS knock-out MNs, and that polysomal localisation of FUS and FMRP is impaired in both knock-out and mutant FUS neurons, we conclude that, albeit interesting, these alterations cannot explain the translation phenotype observed in Fig. 1.

### 3. FUS mutations induce the formation of FMRP condensates

In addition to binding to FUS, FMRP has also been detected in FUS-positive inclusions in *post mortem* tissue (Blokhuis et al., 2016), prompting us to test whether physiological expression of mutant FUS could alter the localisation of this RBP. We focused on MN axons where FMRP condensates are typically distinct. The density of axonal FMRP puncta was increased in a mutant FUS dosage-dependent manner in primary *Fus*^Δ14/+^ and *Fus*^Δ14/Δ14^ MNs (Fig. 2A,B; SFig. 4B; normalised axonal FMRP puncta density in *Fus*^+/+^=100±5.2, *Fus*^Δ14/+^=154.2±8.2, *Fus*^Δ14/Δ14^=181.5±9.2, ****p<0.0001), despite no changes in the somatic intensity of FMRP staining, nor in the total FMRP expression in MN cultures (Fig. 2E; SFig. 4D-F; somatic FMRP fluorescence intensity in *Fus*^+/+^=1±0.04, *Fus*^Δ14/+^=1.2±0.05, *Fus*^Δ14/Δ14^=1.0±0.07; FMRP levels in cultured MNs lysates *Fus*^+/+^=1.9±0.45, *Fus*^Δ14/+^=1.9±0.4, *Fus*^Δ14/Δ14^=1.7±0.4). Puncta size was also unaffected by mutant FUS expression (SFig. 4C; average FMRP axonal puncta size in *Fus*^+/+^=0.26±0.01 μm^2^, *Fus*^Δ14/+^=0.28±0.01 μm^2^, *Fus*^Δ14/Δ14^=0.29±0.01 μm^2^).

**Fig. 2.**
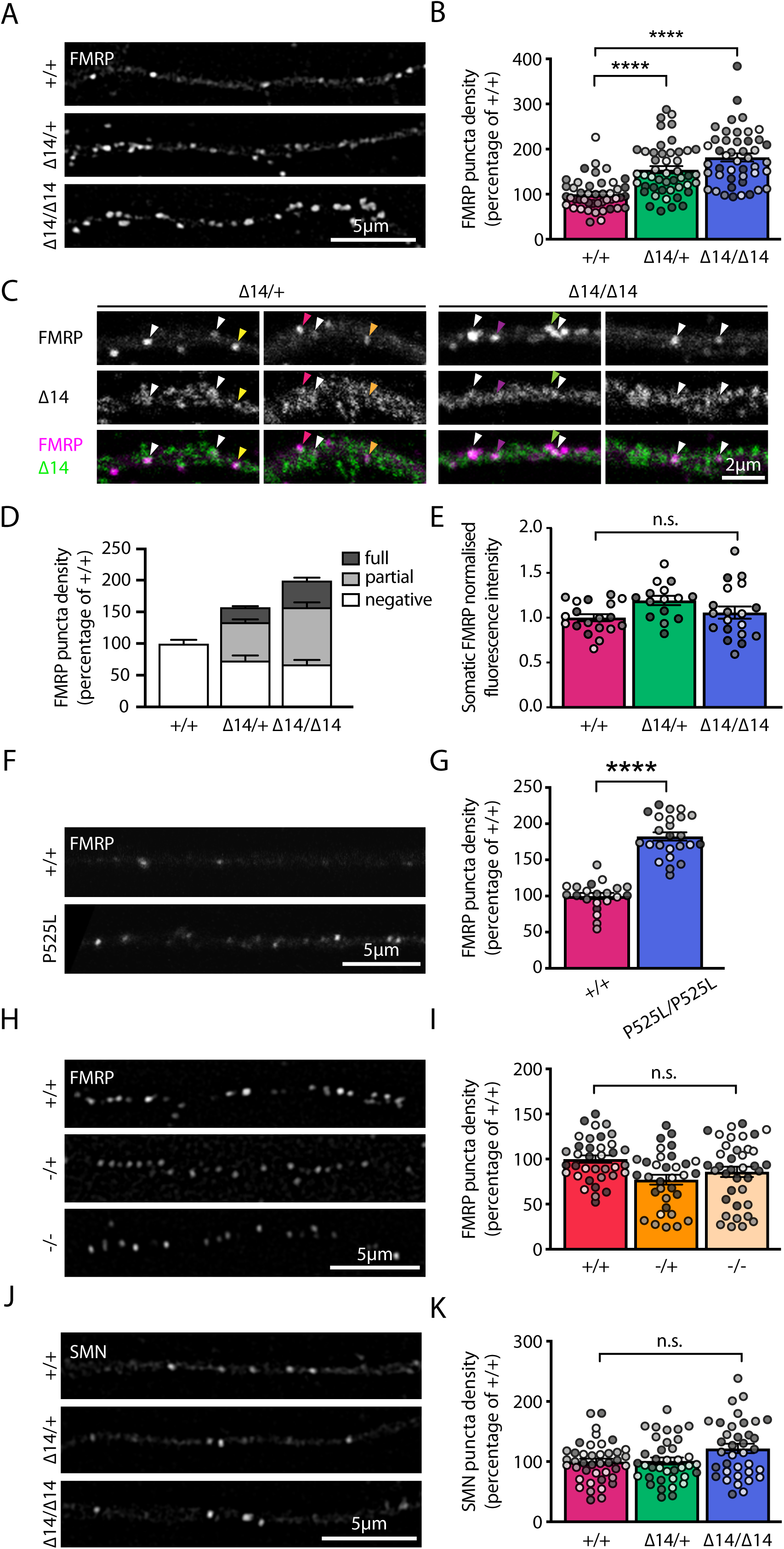
FMRP puncta density is increased in mutant FUS expressing MNs. **(A)** Representative deconvolved images of FMRP axonal puncta in *Fus*^+/+^, *Fus*^Δ14/+^ and *Fus*^Δ14/Δ14^ MNs grown in MFCs (DIV 8). **(B)** Quantification of axonal FMRP puncta density in *Fus*^+/+^, *Fus*^Δ14/+^ and *Fus*^Δ14/Δ14^ MNs (n=4, axons=45-47, ****p<0.0001 Kruskal-Wallis followed by Dunn’s post hoc test) **(C)** Representative images showing axonal FMRP puncta either fully (white arrowheads) or partially (coloured arrowheads) positive for Δ14 FUS in *Fus*^Δ14/+^ and *Fus*^Δ14/Δ14^ MNs. **(D)** Segmentation of FMRP puncta density into fully Δ14 positive, partially Δ14 positive and negative. **(E)** Quantification of somatic FMRP fluorescence in *Fus*^+/+^, *Fus*^Δ14/+^ and *Fus*^Δ14/Δ14^ HB9:GFP positive MNs (n=4, MNs=15=19). **(F)** Representative images of FMRP axonal puncta in *FUS*^+/+^ and *FUS*^P525L/P525L^ iPSC-derived MNs grown in MFCs. **(G)** Quantification of axonal FMRP puncta density in *FUS*^+/+^and *FUS*^P525L/P525L^ iPSC-derived MNs as shown in (F) (n=4, axons=21-24; ****p<0.0001, Student’s t-test). **(H)** Representative deconvolved images of FMRP axonal puncta in *Fus*^+/+^, *Fus*^-/+^ and *Fus*^-/-^ MNs grown in MFCs (DIV 8). **(I)** Quantification of axonal FMRP puncta density in *Fus*^+/+^, *Fus*^-/+^ and *Fus*^-/-^ MNs as shown in (H) (n=3, axons= 33-38). **(J)** Representative deconvolved images of SMN axonal puncta in *Fus*^+/+^, *Fus*^Δ14/+^ and *Fus*^Δ14/Δ14^ MNs grown in MFCs (DIV 8). **(K)** Quantification of axonal SMN puncta density in *Fus*^+/+^, *Fus*^Δ14/+^ and *Fus*^Δ14/Δ14^ MNs (n=3, axons=36-41). Independent experiments are visualised in different shades of grey in the graphs.

Phenocopying the mouse primary motor neurons, we also observed an 80% increase in FMRP axonal puncta in human iPSC-derived *FUS*^P525L/P525L^ MNs (Fig. 2F,G; normalised axonal FMRP puncta density in isogenic control=100±4.4, *FUS*^P525L/P525L^=182.5±5.8, ****p<0.0001). In contrast, we found no alterations in FMRP puncta density in FUS knock-out axons (Fig. 2H,I; normalised axonal FMRP puncta density in *Fus*^+/+^=100±4.1, *Fus*^+/-^=77.26±5.5, *Fus*^-/-^ axons=85.9±5.7), suggesting that an increase in axonal FMRP puncta is caused by a gain of function of mutant FUS. In support of a primary role of FUS in altering FMRP dynamics, ∼60% of FMRP puncta were either fully or partially positive for mutant FUS (Fig. 2C,D; percentage of Δ14 FUS positive FMRP puncta: *Fus*^Δ14/+^ axons 16.3% full and 39.1% partial, *Fus*^Δ14/Δ14^ axons 20.6% full and 45.4% partial).

To assess whether a general impairment in axonal transport could account for the altered FMRP distribution, we analysed the density of LAMP1-positive organelles in MN axons. Mutant FUS expression did not affect axonal density of these structures (SFig. 4G,H; normalised axonal LAMP1 puncta density in *Fus*^+/+^=100±6.5, *Fus*^Δ14/+^=112.6±6.22, *Fus*^Δ14/Δ14^=123.2±8.1), implying that axonal transport is likely unaffected in Δ14 FUS MNs, in agreement with recently published data (Sleigh et al., 2020).

Since we and others have shown that SMN also interacts with FUS (SFig. 4A), and SMN dynamics, particularly in nuclear gems, are affected by ALS-associated FUS mutants (Groen et al., 2013; Scekic-Zahirovic et al., 2016; Sun et al., 2015; Yamazaki et al., 2012), we assessed whether mutant FUS expression affects SMN distribution along axons. We found that SMN puncta density was unaltered in *Fus*^Δ14/+^ and *Fus*^Δ14/Δ14^ MN axons compared to *Fus*^+/+^ axons (Fig. 2J,K; normalised axonal SMN puncta density in *Fus*^+/+^=100±5.2, *Fus*^Δ14/+^=101.1±5.9, *Fus*^Δ14/Δ14^=122.0±7.9), suggesting that only a subset of FUS-binding RBPs, such as FMRP, are affected by mutant FUS.

### 4. FUS mutants induce FMRP incorporation into cytoplasmic condensates by LLPS

To further test the causality of FUS driving FMRP sequestration, we took advantage of the fact that mutant FUS overexpression is known to generate intracellular FUS condensates (Murakami et al., 2015). Overexpression of either NLS-lacking FUS (^mCherry^FUS^513x^) or mutant FUS (^Flag^FUS^P525L^) in HeLa cells induced the formation of large, cytoplasmic FUS-positive condensates (Fig. 3A; SFig. 5A; left panels), which led to a sequestration of endogenous FMRP (Fig. 3A; SFig. 5A). Interestingly, we found that brief (∼18h) overexpression of ^mCherry^FUS^513x^ led to the presence of small FMRP puncta decorating larger FUS condensates (Fig. 3A; top panels), as well as double positive condensates (Fig. 3A; bottom panels), possibly reflecting different phases of incorporation, or a multiphasic behaviour reminiscent of FMRP-Caprin1 condensates (Kim et al., 2019). Overexpression of ^Flag^FUS^P525L^ in primary motor neurons also led to the formation of distinct FUS condensates that were positive for endogenous FMRP (Fig. 3B; arrowheads indicate FMRP-positive FUS puncta).

**Fig. 3.**
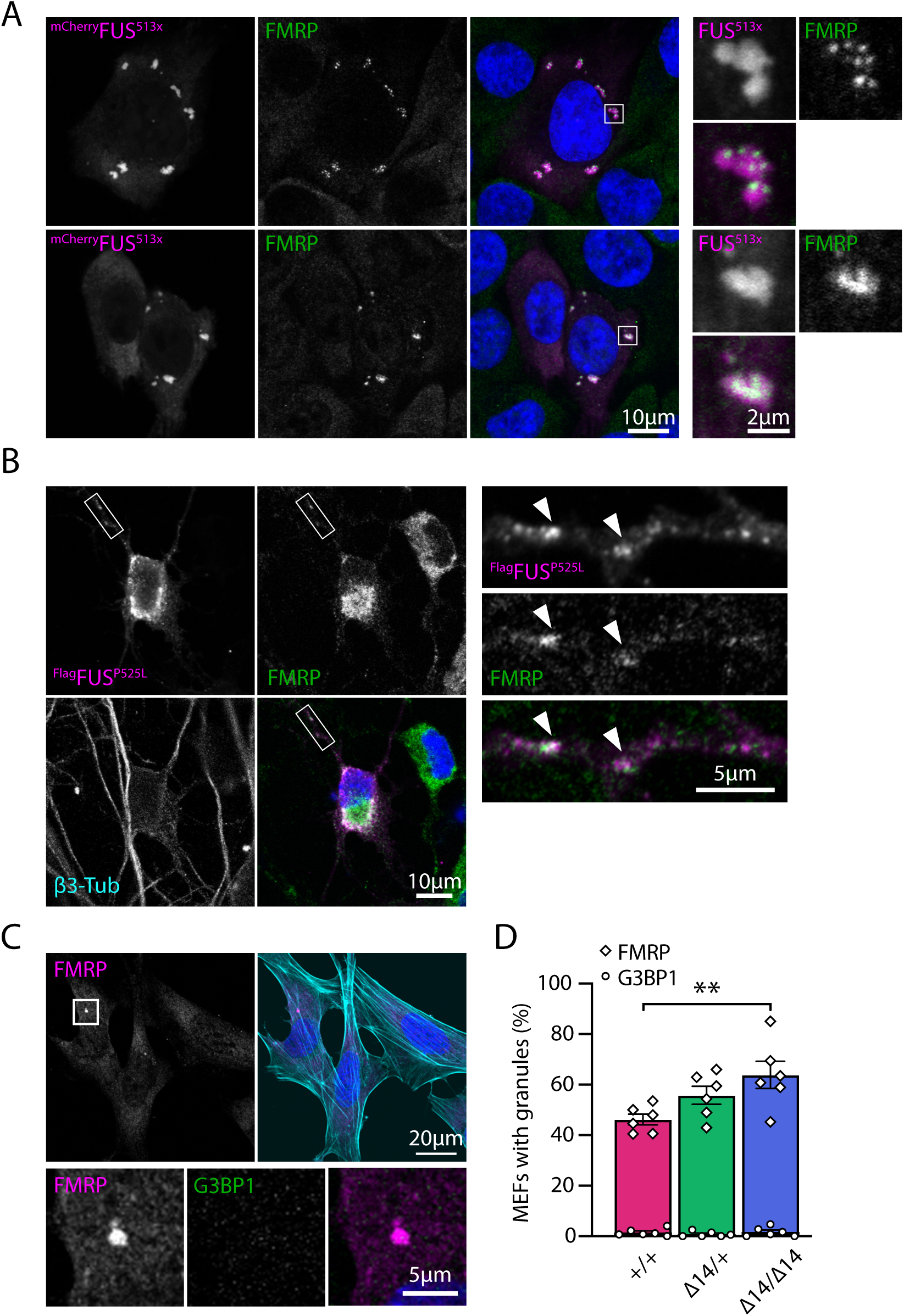
Mutant FUS promotes FMRP incorporation into cytoplasmic condensates. **(A)** Examples of HeLa cells overexpressing ^mcherry^FUS^513x^. Endogenous FMRP (middle panels, green) is recruited to ^mcherry^FUS^513x^ condensates and either forms puncta that decorate the condensates (top panels) or partitions into them (bottom panels). On the right, zoomed images of the condensates in the merged images white boxes. **(B)** Overexpression of ^Flag^FUS^P525L^ (magenta) in primary MNs forms discrete puncta that are positive for endogenous FMRP (green). On the right, enlargement of the neurite in the white box, arrowheads indicate FMRP positive ^Flag^FUS^P525L^ puncta. **(C)** Representative images of typical FMRP condensates in MEF cells (*Fus*^Δ14/+^). In the bottom panels zoomed images of the white box showing that spontaneous FMRP condensates (magenta) are negative for the stress granule marker G3BP1 (green). **(D)** Quantification of the percentage of *Fus*^+/+^, *Fus*^Δ14/+^ and *Fus*^Δ14/Δ14^ MEF cells with FMRP puncta (>0.5mm^2^, as shown in (C) - diamond shape) and G3BP1 puncta (circle) (n=6, **p<0.01; paired Friedman test followed by Dunn’s multiple comparisons post hoc test).

Since FUS overexpression may induce stress granules, FMRP could be recruited in response to cellular stress, rather than as a direct consequence of the presence of cytoplasmic mutant FUS. To address this, we generated mouse embryonic fibroblasts (MEFs) from our Δ14 FUS mouse model, which expresses mutant FUS at physiological levels. When we looked at endogenous FMRP, we found that the presence of mutant FUS led to a dose-dependent increase in the number of cells with spontaneous FMRP condensates (>0.5 μm^2^) compared to control (Fig. 3C,D; SFig. 5B; percentage of MEFs with FMRP puncta: *Fus*^+/+^=46.2±2.1%, *Fus*^Δ14/+^=55.8±3.6%, *Fus*^Δ14/Δ14^=63.8±5.4%; **p<0.05). This did not coincide with alterations in the number of condensates per cell (SFig. 5C; average puncta number per cell in *Fus*^+/+^ 2.7±0.2; *Fus*^Δ14/+^ 2.3±0.2; *Fus*^Δ14/Δ14^ 3.5±0.3) or condensate size (SFig. 5D), and the large majority of these structures were negative for stress granule markers, such as G3BP1 (Fig. 3C,D; SFig. 5B; percentage of MEFs with G3BP1 puncta *Fus*^+/+^=1.5±0.6%, *Fus*^Δ14/+^=0.8±0.4%, *Fus*^Δ14/Δ14^=1.7±0.7%). Together, this further supports that cytoplasmic mutant FUS regulates the dynamics and localisation of FMRP in neuronal and non-neuronal cells, and that the presence of FMRP in these structures is not just secondary to the formation of stress granules.

To further explore FUS-induced FMRP behaviour, we asked whether the two proteins could co-phase separate *in vitro*. We performed an *in vitro* co-LLPS assay in which the disordered N-terminal region of FUS conjugated with FITC (^FITC^FUS_LCD_) was co-incubated with the disordered C-terminal region of FMRP conjugated with AlexaFluor-647 (^Alexa-647^FMRP_LCD_), in the presence of sc1, a G-quadruplex-forming RNA known to bind FMRP (Phan et al., 2011; Tsang et al., 2019). We observed droplets that were positive for both ^FITC^FUS_LCD_ and ^Alexa-647^FMRP_LCD_ (Fig. 4A), evidence that the FUS disordered N-terminal region interacts with sc1 RNA and FMRP.

**Fig. 4.**
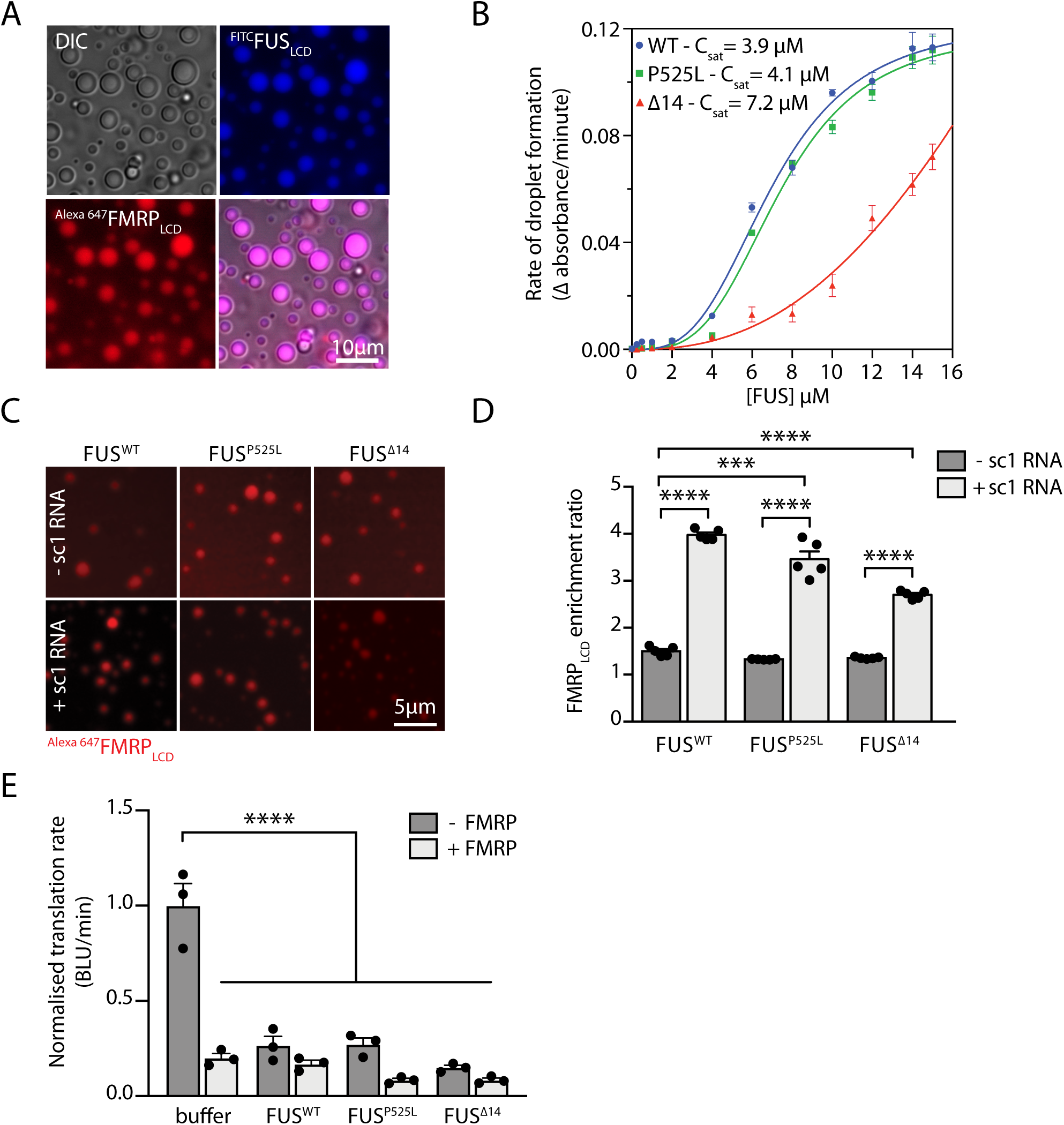
Recombinant FUS promotes FMRP LLPS and inhibits translation *in vitro*. **(A)** *In vitro* co-LLPS assay of ^FITC^FUS^LCD^ with ^Alexa647^FMRP^LCD^ in the presence of sc1 RNA. 50 μM of each protein and 1 μM of sc1 RNA were used (n=3). **(B)** Phase separation propensities of different FUS constructs are determined by the change in turbidity as a function of time. Each point represents the mean rate of turbidity change and error bars represent the SEM (n=3). **(C)** Representative images of an *in vitro* co-LLPS assay showing the partitioning of ^Alexa-647^FMRP^LCD^ into wild type, P525L or Δ14FUS mutants in the presence or absence of sc1 RNA. FUS (10 μM) phase separation was induced by TEV protease (0.5 μM) cleavage before the addition of ^Alexa647^FMRP^LCD^ (1 μM) and sc1 RNA (0.5 μM). Scale bar represents 5 μm. **(D)** Quantification of ^Alexa647^FMRP^LCD^ enrichment into FUS^WT^, FUS^P525L^ and FUS^Δ14^ droplets as shown in (C) (n=5; ***p<0.001, ****p<0.0001 one way ANOVA followed by Tukey’s multiple comparison test). **(E)** Recombinant FUS^WT^, FUS^P525L^ and FUS^Δ14^ (10 μM) phase separation was induced by TEV protease (0.5 μM) cleavage, and proteins added to an *in vitro* rabbit reticulocyte translation system with luciferase mRNA in absence or presence of FMRP (10 μM). Change in bioluminescence (BLU) rate is used as a reporter for translational activity. All results were normalized to buffer control (+ TEV) (n=3; ****p<0.0001, one way ANOVA followed by Sidak multiple comparison test).

To analyse the partitioning of FMRP into droplets of full-length FUS, we generated wildtype, P525L and Δ14 recombinant FUS, and to circumvent the strong propensity of FUS to aggregate, an MBP tag was added to FUS to enhance solubility (SFig. 6A). Upon cleavage of the MBP tag by TEV protease, all FUS proteins phase separated *in vitro* as detected by turbidity measurements (Fig. 4B). FUS^Δ14^ displayed a lower LLPS propensity compared to FUS^WT^ and FUS^P525L^, likely due to the loss of the C-terminal RG/RGG region (SFig. 6C). To investigate FMRP incorporation into FUS condensates, ^Alexa-647^FMRP_LCD_ was added to pre-formed FUS droplets in the presence or absence of sc1 RNA. In the absence of sc1 RNA, FMRP was enriched equally in FUS^WT^, FUS^P525L^ and FUS^Δ14^ condensates, however, in the presence of sc1 RNA, the enrichment of ^Alexa-647^FMRP_LCD_ into these structures was increased for all FUS proteins (Fig. 4C,D; ^Alexa-647^FMRP_LCD_ enrichment ratio in FUS^WT^=1.5±0.04, FUS^WT^+sc1=4.0±0.05, FUS^P525L^=1.3±0.004 FUS^P525L^+sc1=3.5±0.17, FUS^Δ14^=1.4±0.008, FUS^Δ14^+sc1=2.7±0.04; ***p<0.001, ****p<0.0001). FUS^Δ14^ droplets have a lower FMRP enrichment compared to those formed by FUS^WT^ and FUS^P525L^, possibly due to the loss of the RG/RGG motifs that could lead to decreased critical RG/RGG-RNA interactions and therefore reduce FMRP partitioning. These results show that FMRP partitions into FUS-droplets and that this process occurs via protein-protein and protein-RNA interactions.

### 5. FUS and FMRP repress translation *in vitro*

Phase separated condensates are considered to represent a state antagonistic to translation (Krichevsky and Kosik, 2001; Sahoo et al., 2018), and FMRP phase separation correlates with translation inhibition (Tsang et al., 2019). To test whether FUS condensates can directly affect protein synthesis we took advantage of an *in vitro* translation assay, in which recombinant MBP-FUS fusion proteins were directly added to a commercial rabbit reticulocyte cytoplasmic extract and its phase separation was induced by TEV protease cleavage. Quantification of the bioluminescence of the firefly luciferase, as a reporter for the translation of its mRNA, showed that all FUS proteins significantly suppressed translation, and addition of FMRP_LCD_ induced a further decrease in luciferase translation (Fig. 4E; normalised bioluminescence (BLU)/minute in buffer control=1±0.12, buffer+FMRP=0.2±0.02, FUS^WT^=0.27±0.05, FUS^WT^+FMRP=0.17±0.02, FUS^P525L^=0.27±0.03, FUS^P525L^+FMRP=0.08±0.01, FUS^Δ14^=0.15±0.01, FUS^Δ14^+FMRP=0.08±0.01; ****p<0.0001). These results support FUS having a repressive role on protein synthesis and that mislocalisation of the protein to the cytoplasm is key to driving this gain of function mechanism. In addition, FMRP has an additive effect on translation inhibition.

### 6. Mutant FUS impairs translation of FMRP target RNAs *in vivo*

We next investigated whether mutant FUS expression, and the consequent dysregulation of FMRP dynamics, could affect *in vivo* the translation of RNAs that specifically bind FUS and FMRP. We crossed either our FUS mutant or FUS knock-out lines with mice expressing both the Cre-dependent HA-tagged RPL22 ribosomal subunit (*Rpl22^HA^*, RiboTag) and a MN-specific Cre-recombinase (*Chat-Cre*) (Sanz et al., 2009), allowing us to immuno-purify transcripts bound by ribosomes in MNs, referred to as the “translatome” throughout the manuscript (Fig. 5A; SFig. 7A-C). RNA pulldown was specific to the presence of the Cre-recombinase (SFig. 7A,B), and qPCR showed enrichment for the motor neuronal genes *Chat* and *Rpl22^HA^*, and depletion of glial gene *Pmp22*, confirming MN-specificity of pulldown (Fig. 5B).

**Figure 5.**
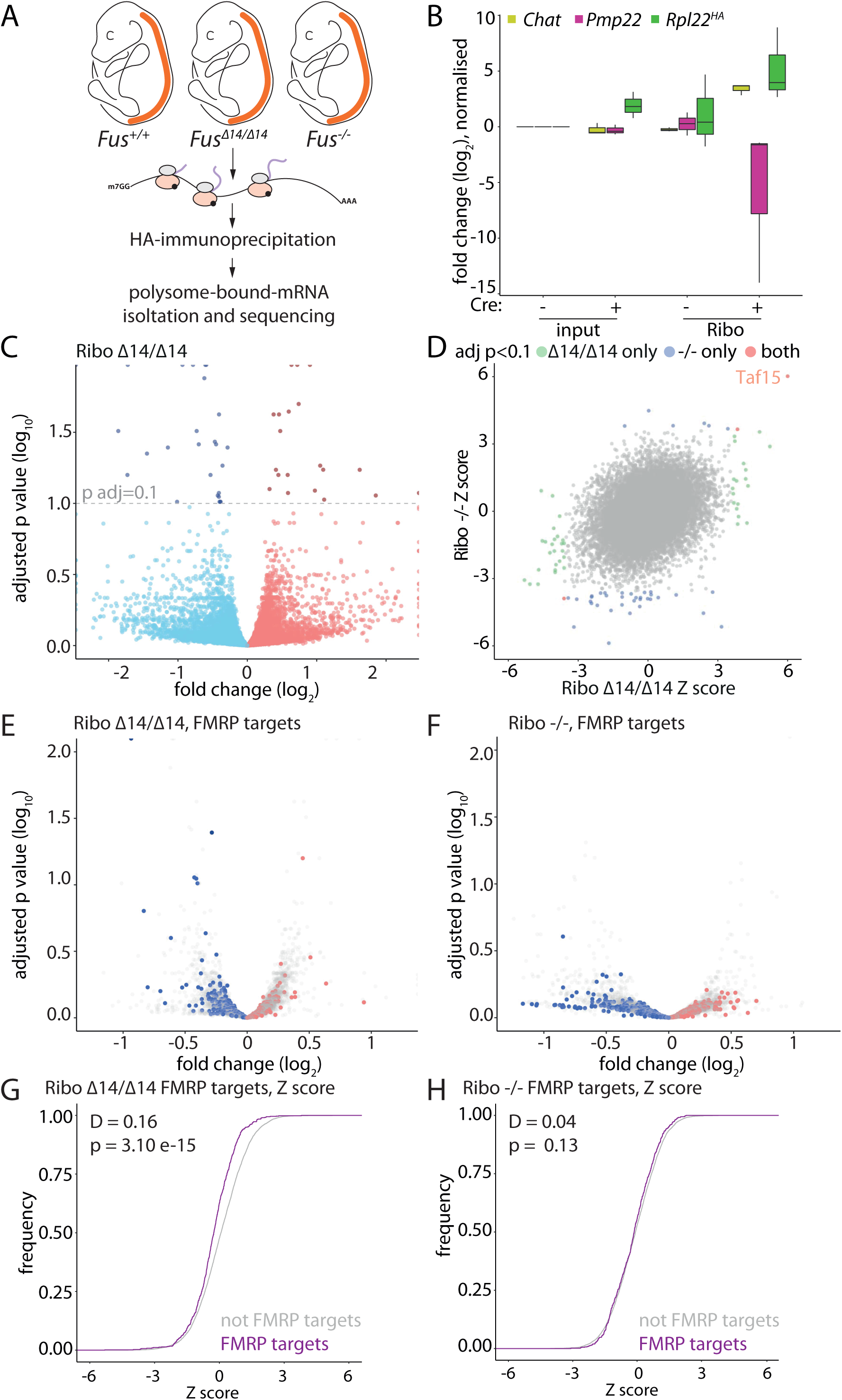
Translation of FMRP-bound genes is decreased in motor neurons *in vivo*. **(A)** RiboTag method was used to purify MN-specific, translation-engaged transcripts from embryonic spinal cords (E17.5) of *Fus^Δ14/Δ14^* and *Fus^-/-^* and their wildtype littermates. **(B)** qPCR analysis of total spinal cord tissue (input) and HA-tagged, ribosome-associated, MN-specific fraction (Ribo). Expression of MN markers: *Chat* and *Rpl22^HA^* and the glial marker *Pmp22* was measured, *Gapdh* expression was used as housekeeping control (adult spinal cord tissue, 3 months of age, n=3). **(C)** Volcano plot of MN-specific translatome Ribo-*Fus^Δ14/Δ14^*. Blue points: fold change (log_2_) < 0; red points: fold change (log_2_) > 0, genes with fold change (log_2_) > 2.25 or < −2.25 or with adjusted p value (-log_10_) < 2 were plotted as infinity, (n=5). **(D)** Distribution of Z scores in the MN-specific translatome Ribo-*Fus^Δ14/Δ14^* and Ribo-*Fus^-/-^* shows mutation-specific changes. Green points: adjusted p value < 0.1 for *Fus^Δ14/Δ14^* only, blue points: adjusted p value < 0.1 for *Fus^-/-^* only, red points: adjusted p value < 0.1 for *Fus^Δ14/Δ14^* and *Fus^-/-^*. **(E)** Volcano plot of MN-specific translatome Ribo-*Fus^Δ14/Δ14^* (filtered by expression (log_10_) base mean < 4.25 and > 2.5, FMRP targets in red and blue, not FMRP targets in grey). **(F)** Volcano plot of MN-specific translatome Ribo-*Fus^-/-^* filtered by expression (log_10_) base mean < 4.25 and > 2.5, FMRP targets in red and blue, no FMRP targets in grey. **(G)** Cumulative frequency plot of Z scores of genes in (E) shows a significant decrease of FMRP targets expression in MN-specific, translatome of *Fus^Δ14/Δ14^* (Kolmogorov-Smirnov test) **(H)** Cumulative frequency plot of Z scores of genes shown in (F) shows no change of FMRP targets expression in MN-specific, translatome of *Fus^-/-^* (Kolmogorov-Smirnov test).

Importantly, FUS mutations, as well as loss of FUS, induce alterations in transcript expression, with the latter having a stronger effect (Humphrey et al., 2020). These changes can influence the MN translatome composition, nonetheless we found alterations in our translatome datasets that did not correlate with a change in the total spinal cord RNA (input) expression, both in this dataset, and in our previous high-depth RNA-seq data (Humphrey et al., 2020) (**Supplementary Table 1**). Further, as MN-derived transcripts are a minority of the total spinal cord RNA, we set out to understand whether mutant FUS had an additional gain of function on the MN translatome by comparing the effect of mutant FUS expression to that of FUS knock-out. We performed differential expression analysis of the *Chat-Cre*/*Fus*^Δ14/Δ14^/*Rpl22^HA^* translatome, henceforth referred to as Ribo*-Fus*^Δ14/Δ14^ (Fig. 5C; **Supplementary Table 1**; 21 upregulated and 26 downregulated transcripts, adjusted p value < 0.1) and the *Chat-Cre*/*Fus*^-/-^ /*Rpl22^HA^* translatome, referred to as Ribo-*Fus*^-/-^ (SFig. 7D; **Supplementary Table 1**; 8 upregulated and 39 downregulated transcripts, adjusted p value < 0.1) compared to their own littermate wildtype controls. We calculated a Z score for each gene in the Ribo-*Fus*^Δ14/Δ14^ and Ribo-*Fus*^-/-^ experiments and found a significant correlation between transcripts that were altered within both datasets (Pearson correlation coefficient *r*=0.28, p value < 2.2e-16). We identified changes that were common between Ribo-*Fus*^Δ14/Δ14^ and Ribo-*Fus*^-/-^ and others that were specific to the translatome of each genotype (Fig. 5D; **Supplementary Table 2**; 3 transcripts with an adjusted p value < 0.1 in both Ribo*-Fus*^Δ14/Δ14^ and Ribo*-Fus*^-/-^, 43 transcripts only in Ribo*-Fus*^Δ14/Δ14^ and 39 only in Ribo*-Fus*^-/-^). Our results highlight that *Taf15*, one of the three members of the FET family (along with *Fus* and *Ewsr1*), is increased in both Ribo-*Fus*^Δ14/Δ14^ and Ribo-*Fus*^-/-^ datasets. Moreover, previous findings have shown that mutant FUS impairs its autoregulation leading to an increase in RNA levels (Humphrey et al., 2020); we here find that *Fus* is increased in the mutant translatome dataset (Fig. 5D), indicating that impaired autoregulation could indeed alter FUS protein levels.

We next asked whether there were changes in ribosome-association of RNAs bound by FUS and FMRP, identified using published and widely-used CLIP data (Darnell et al., 2011; Rogelj et al., 2012). As FUS, unlike FMRP, mostly binds pre-mRNA intronic sequences, we selected only transcripts where FUS binds within the mature RNA sequence (5ʹ UTR, exons and 3ʹ UTR). To avoid any bias deriving from differences in expression levels, we compared target RNAs to a set of non-target transcripts with similar expression (SFig. 8 for FUS; SFig. 9 for FMRP). We plotted the distribution of target and non target RNAs (SFig. 8B) and, as in previous translatome analysis, we compared their cumulative frequency of Z scores, with a shift of the curve towards the right indicating a general upregulation of targets in the condition of interest, and a leftward shift indicating a overall downregulation (Goering et al.; Thomson et al., 2017). The comparison between FUS targets versus non target controls shows a minor, albeit significant, right shift of the cumulative distribution in Ribo-*Fus*^Δ14/Δ14^ (distance (D)=0.05; p=1.20 e-05), while no change was detected in the Ribo-*Fus*^-/-^ dataset (SFig. 8C; D=0.02; p=0.26). Conversely, when investigating FMRP targets, we found more widespread changes (Fig. 5E,F; SFig. 9B), with greater distances even in the total spinal cord samples (input), (SFig. 9C; *Fus*^Δ14/Δ14^ left shift, D=0.16, p value=2.56 e-14; *Fus*^-/-^ left shift, D=0.24, p value=2.2e-16). Interestingly, however, when we analysed the translatome datasets, we found that FMRP target distribution was altered selectively in Ribo-*Fus*^Δ14/Δ14^ (Fig. 5G; left shift, D=0.16, p value=3.10 e-15), while no distribution change was present in Ribo-*Fus*^-/-^ (Fig. 5H; D=0.04, p value=0.13). This indicates that in addition to alterations in the expression, mutant FUS leads to impaired ribosome-association of FMRP targets.

## Discussion

Cytoplasmic mislocalisation and nuclear depletion of FUS are hallmarks of FUS-ALS, and the degree of cytoplasmic misplacement induced by different disease-causing mutations correlates with disease severity (Dormann et al., 2010). FUS expression levels, similarly to many RBPs, are physiologically highly regulated and both exogenous overexpression and complete depletion were found to impact on numerous cellular processes, causing lethality (Mitchell et al., 2013; Qiu et al., 2014; Scekic-Zahirovic et al., 2016; Sephton et al., 2014). Only with the generation of physiological models of FUS-ALS has it become clear that, in addition to a nuclear loss of function, a cytoplasmic gain of function is necessary to drive cellular toxicity and ultimately MN loss (Scekic-Zahirovic et al., 2016). However, the molecular mechanisms underlying this process are unclear and alterations in protein translation and in cytoplasmic granule formation have been hypothesised. Here, we have used both mouse and human models with endogenous expression of FUS mutants to establish that ALS-causing FUS mutations repress translation (Fig. 1C,D,F,G). Although we found that mutant FUS association with polysomes is impaired, this is not sufficient to induce the translation deficit. In addition, we show that, both *in vitro* and *in vivo*, FUS forms condensates where FMRP, another RBP strongly linked to neurodegeneration and translation (Cid-Samper et al., 2018; Richter et al., 2015; Tan et al., 2020; Todd et al., 2013), is sequestered. Finally, we demonstrate that translation of FMRP target RNAs is impaired *in vivo*, establishing a pathogenesis paradigm by which FUS impairs translation by altering the dynamics of other RBPs (Fig. 6).

**Fig. 6.**
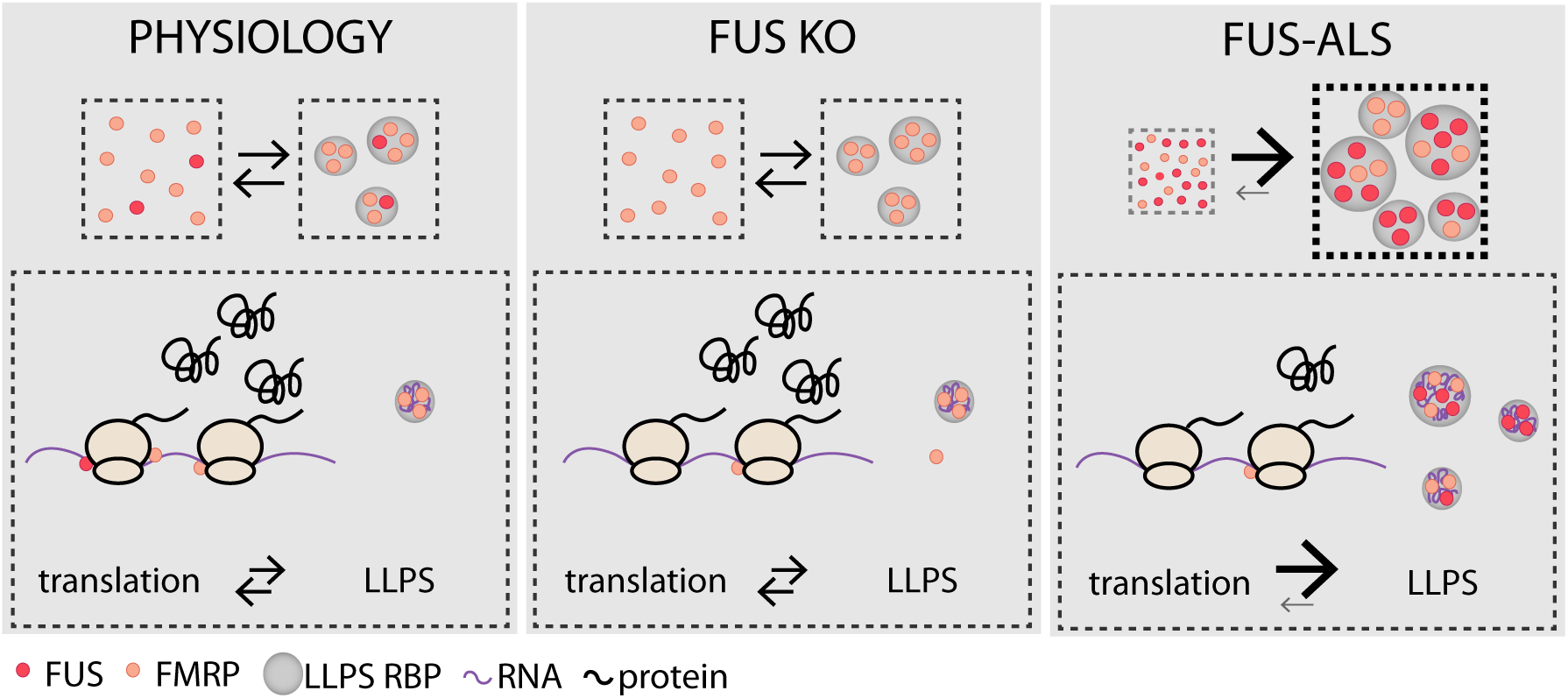
Proposed model of mutant FUS cytoplasmic gain of function. In control conditions (left panel) low levels of FUS are present in the cytoplasm and the phase-separation of FUS and FMRP are at a physiological equilibrium. Loss of FUS (middle panel) results in a reduction of FMRP association with the translational machinery, however this does not induce significant alterations in FMRP LLPS or global protein translation. In FUS-ALS (right panel), the increased cytoplasmic localisation of FUS shifts the LLPS equilibrium of both FUS and FMRP resulting in an increase in cytoplasmic condensates. This is associated with a depletion of the proteins from the translational machinery and an overall decrease in protein synthesis.

Recently, numerous studies have demonstrated that FUS can undergo LLPS and that cytoplasmic FUS granules are, indeed, phase separated condensates (Murakami et al., 2015; Patel et al., 2015). Interestingly, our *in vitro* experiments show that, compared to FUS^WT^ and FUS^P525L^, FUS^Δ14^ has a lower propensity to undergo LLPS (Fig. 4B). This is likely due to the lack of the C-terminal RGG/RG region (SFig. 6C) and highlights how a combination of mislocalisation and LLPS dynamics can determine FUS cytoplasmic gain of function. In fact, NLS-lacking FUS mutants have a stronger mislocalisation compared to NLS missense mutants, such as FUS^P525L^; however, the stronger LLPS propensity of the latter may result in comparable cellular toxicity.

Mutant Δ14 FUS condensates are present throughout the MN cell body and neurites (Fig. 1A,B). Since phase separated condensates, such as transport granules or stress granules, are macromolecular complexes typically in a translationally inactive state, one possibility is that the presence of these FUS condensates could indirectly impact on translation. Further supporting this hypothesis, disassembly of RNA granules is associated with protein translation. For example neuronal activity triggers β-actin granule disassembly, associated with activity-dependent protein synthesis (Buxbaum et al., 2014; Park et al., 2014); and similarly, injury-induced dissolution of G3BP1 granules results in the translation of G3BP1-bound RNAs (Sahoo et al., 2018).

Within cells, the composition of phase separated condensates is heterogeneous and, while LLPS is common between LCD-containing proteins, *in vitro* studies have demonstrated that FUS is particularly prone to undergo this process. Moreover, FUS phase separation can favour the co-partitioning of so-called ‘client proteins’, that contain LCDs, but would not normally form condensates at physiological concentrations (Wang et al., 2018).

With this in mind, we questioned whether the presence of cytoplasmic FUS condensates could alter the distribution and dynamics of a wider RBP network. Since some FUS-interacting RBPs are also found in FUS inclusions in *post mortem* tissue or are disrupted in models of FUS-ALS (Blokhuis et al., 2013, 2016; Scekic-Zahirovic et al., 2016), we decided to analyse the distribution of the two most well characterised ones: FMRP and SMN. When we analysed the distribution of axonal FMRP, we found a dose-dependent increase in FMRP condensates in mutant FUS-expressing MNs (Fig. 2A,B,F,G), a change that we observed with two ALS-causing FUS mutations, that is not dependent on FMRP expression levels and is conserved in iPSC-derived MNs. We investigated the axonal distribution of these RBPs since they have key roles in RNA transport and axonal function. Axonal analysis also allowed us to detect and analyse distinct puncta, however, it is likely that the effect is not restricted to this compartment, in fact we found alterations in the frequency of FMRP condensates also in non-neuronal MEF cells (Fig. 3D; SFig. 5B). Although FMRP granules can be induced by cellular stress, FUS-induced FMRP condensates are negative for stress granule markers (Fig. 3C,D; SFig. 5B); we therefore propose that the increased FMRP condensate formation is due to a cytoplasmic gain of function of mutant FUS.

Interestingly, we did not find any alteration in the distribution of SMN (Fig. 2J,K). While the expression of mutant FUS may still affect SMN functionality (for example at nuclear level, as previously reported (Scekic-Zahirovic et al., 2016)), this indicates that, in addition to protein-protein interaction, other intrinsic or extrinsic factors are required to promote FUS-induced condensate formation.

While FUS-mediated translation regulation has not yet been studied in detail, the role of FMRP as a translation repressor is well established. FMRP inhibits translation through several mechanisms, including polysome-binding, miRNA and RISC-dependent repression, or by sequestering translation initiation factors (Richter et al., 2015). More recently, FMRP LLPS has been shown to correlate with translation inhibition *in vitro* (Tsang et al., 2019), adding complexity to the FMRP-dependent regulation of translation, and supporting a model in which an increase in FMRP condensates could be associated with decreased translation of FMRP targets. To explore how the translation landscape is affected by mutant FUS expression, we have sequenced ribosome-engaged transcripts from motor neurons. Importantly, FUS mutations induce transcript expression changes, and we have previously shown that these are comparable to, but weaker than FUS knock-out (Humphrey et al., 2020). However, when we compared the translatome datasets, we found specific changes in the Δ14 FUS compared to the knock-out translatome (Fig.5 D), indicating that mutant FUS affects the translatome through a gain of function mechanism.

Within the upregulated transcripts we have also identified FUS itself. This is in agreement with mutant FUS being defective at autoregulating its own transcript levels (Humphrey et al., 2020; Zhou et al., 2013), and further supports that the altered transcript expression may indeed result in increased protein translation, which, in turn, could worsen its cytoplasmic partitioning and create a vicious cycle leading to increased FUS and FMRP condensate formation.

When we analysed FMRP target RNAs, we found that both mutant FUS expression and FUS depletion result in altered transcript expression levels. However, when we looked at the distribution of FMRP target RNAs in our translatome data, we found a selective depletion of these transcripts from the purified ribosomal fractions of our mutant Δ14 FUS model (Fig.5 G). This depletion is strikingly similar to that described in a FMRP knock-out model (SFig.9 C; re-analysed from (Thomson et al., 2017) Ribo-*Fmr1*^-/y^ D=0.18, p value=2.56 e-11; total RNA *Fmr1*^-/y^ D=0.19, p value<2.2 e-16).

Rather than a direct role on polysomal function, our data supports a model in which increased FMRP condensate formation, induced by mutant FUS, reduces the regulation and consequent availability of FMRP targets, ultimately altering their translation pattern (Fig. 6) in a way that is comparable to FMRP knock-out. In agreement with FMRP having a concurrent role in FUS-mediated toxicity, FMRP co-expression can rescue FUS-dependent denervation in zebrafish (Blokhuis et al., 2016). However, since FMRP LLPS dynamics, rather than expression levels, are affected by mutant FUS expression, and given that overexpression of FMRP itself promotes its LLPS, we believe it is unlikely that it could rescue translation in our model, although it may have some localised effect on specific targets (Garone et al., 2020b).

Cytoplasmic mislocalization of FUS also occurs in the absence of disease-associated mutations, both in ALS cases caused by other genetic determinants (Tyzack et al., 2019) and in cases of frontotemporal lobar degeneration (FTLD) (Lashley et al., 2011; Neumann et al., 2009). In our *in vitro* translation assays we show that wildtype FUS impairs translation in a similar manner to ALS mutants (Fig. 4E), and although FUS-ALS mutations may also impact on the biophysical properties of the condensates, causing a worsening of the phenotype, FUS mislocalisation may be sufficient to drive translation repression in these pathologies.

Our results support a model whereby the presence of cytoplasmic, phase separated mutant FUS alters the dynamics of a wider network of RBPs. FMRP was found to ameliorate the phenotypes induced by ALS mutations in both FUS and TDP-43 (Blokhuis et al., 2016; Coyne et al., 2015). Moreover, it was recently shown that an altered cross-regulation between FUS, FMRP and the RBP HuD results in an aberrant axonal phenotype in FUS-ALS models (Garone et al., 2020b). It is therefore likely that FUS and FMRP, and possibly other LCD-containing RBPs with a similar biophysical behaviour, form a wider network, and alterations in their cytoplasmic localisation can influence their LLPS dynamics. This would result in the generation of phase separated heterogeneous condensates, in which the RNAs that are bound by RBPs present in the condensates are sequestered and their translation is inhibited. Impaired translation can impact overall neuronal functionality. FUS mislocalisation and the aberrant phase separation of this RBP network can alter these dynamics, possibly contributing to ALS pathogenesis and ultimately affecting motor neuron survival.

## Materials and Methods

### Data Availability

Raw RNA sequencing data from Ribotag input and IP (see below) was deposited at the Gene Expression Omnibus with accession (pending).

### Animals

Δ14 *Fus* mice (B6N;B6J-Fus^tm1Emcf/H^, MGI MGI:6100933) were previously described, (Devoy et al., 2017). *Fus* knock-out mice were obtained from the Mouse Knockout Project (Fustm1(KOMP)Vlcg). HB9::GFP (B6.Cg-Tg(Hlxb9-GFP)1Tmj/J), ChAT-IRES-Cre (B6;129S6-*Chattm2(cre)Lowl*/J) and RiboTag (B6N.129-*Rpl22tm1.1Psam*/J) mice were obtained from the Jackson laboratory. All mouse lines were backcrossed onto C57BL/6J animals for more than 5 generations. Both Δ14 *Fus* knock-in and *Fus* knock-out animals were maintained in heterozygosity, since homozygous mice die perinatally. Both mouse lines were crossed with heterozygous HB9:GFP mice when required.

For RiboTag experiments both RiboTag and ChAT-Cre homozygous mice were crossed with either *Fus^Δ14/+^* or *Fus^+/-^* animals. Double transgenic mice were subsequently crossed to obtain experimental progeny.

All experiments were carried out following the guidelines of the UCL Queen Square Institute of Neurology Genetic Manipulation and Ethics Committees and in accordance with the European Community Council Directive of 24 November 1986 (86/609/EEC). All procedures for the care and treatment of animals were carried out under license from the UK Home Office in accordance with the Animals (Scientific Procedures) Act 1986 Amendment Regulations 2012, and were approved by the UCL Queen Square Institute of Neurology Ethical Review Committee.

### Primary motor neuron preparation

E12.5-14.5 embryos for ventral horn cultures were obtained from heterozygous *Fus^Δ14/+^* or *Fus^-/+^* mice (with or without the HB9:GFP transgene). Briefly, embryos were euthanised, a sample of the tail was used for genotyping and the body maintained in ice cold Hibernate-E media supplemented with B27. Spinal cords of the correct genotype were then dissected, meninges removed and dorsal horns resected. Spinal cord ventral horns were incubated in 0.025% trypsin for 10 min at 37 °C. Trypsin was then removed and the tissue triturated in L15 media containing 0.4% BSA and 0.1 mg/ml DNAse. Neurons were pelleted through a 4% BSA cushion, resuspended in Neurobasal media (Thermo Fisher) containing 2% heat inactivated horse serum, 1x B27, 1x Glutamax, 1x penicillin/streptomycin, 24.8 μM β-mercaptoethanol, 1 ng/ml BDNF, 0.1 ng/ml GDNF, 10 ng/ml CNTF, and immediately plated onto 13mm coverslips, microfluidic chambers or 3 cm dished that had been pre-coated first with 10 μg/mL poly-ornithine and 3 μg/ml laminin. Neurons were maintained in culture in a humidified incubator at 37°C with 5% CO_2_ for 5-7 days in vitro (DIV). Motor neurons were transfected at DIV 2 by magnetofection as previously described (Fallini et al., 2010).

### iPSC maintenance and differentiation

Human iPSCs used in this study are the isogenic FUS^WT/WT^ and FUS^P525L/P525L^ lines were derived and maintained as described (Lenzi et al., 2015), and differentiated into spinal motor neurons as described (De Santis et al., 2019; Garone et al., 2020a). Briefly, iPSCs stably transduced with a piggyBac vector carrying inducible Ngn2, Isl1 and Lhx3 (NIL) transgenes dissociated to single cells with Accutase (Thermo Fisher Scientific) and plated in Nutristem-XF/FF medium (Biological Industries) supplemented with 10 μM rock inhibitor (Enzo Life Sciences) on Matrigel (BD Biosciences) at a density of 100,000 cells/cm^2^. The day after NIL expression was induced by adding 1 μg/ml doxycycline (dox) (Thermo Fisher) in DMEM/F12 medium (DMEM/F12, Sigma Aldrich, supplemented with 1x Glutamax, Thermo Fisher, 1x NEAA, Thermo Fisher, and 0.5x Penicillin/Streptomycin, Sigma Aldrich). The medium was changed every day. After 48h of dox induction, the medium was changed to Neurobasal/B27 medium (Neurobasal Medium, Thermo Fisher, supplemented with 1x B27, Thermo Fisher, 1x Glutamax, Thermo Fisher, 1x NEAA, Thermo Fisher, and 0.5x penicillin/streptomycin, Sigma Aldrich), containing 5 μM DAPT and 4 μM SU5402 (both from Sigma Aldrich). At day 5, cells were dissociated with Accutase (Thermo Fisher) and plated on Matrigel (BD Biosciences) coated dishes or coverslips at the density of 100,000 cells per cm^2^. 10 μM ROCK inhibitor was added for the first 24 h after dissociation. Neuronal cultures were maintained in neuronal medium (Neurobasal/B27 medium supplemented with 20 ng/ml BDNF, 10 ng/ml GDNF, both from PreproTech, and 20 ng/ml L-ascorbic acid, Sigma Aldrich).

### MEFs and cell lines

Mouse embryonic fibroblasts (MEFs) were isolated from the embryonic tissue discarded from the primary motor neuron preparation. Viscera were removed, the remaining tissue was triturated with a blade and incubated in 0.25% trypsin for 20 min at 37°C. Trypsin was quenched and cells were plated in Dulbecco’s modified Eagle media (DMEM, Thermo Fisher) containing 10% fetal bovine serum (FBS) and 1x penicillin/streptomycin. After three days in culture, cells were immortalised by transfection by lipofectamine 3000 (Invitrogen) with the simian virus 40 (SV40) T antigen. After 5–7 passages at low density, the cultures presented a homogeneous cell population and started to grow steadily.

HeLa and MEF cells were maintained in DMEM containing 10% fetal bovine serum (FBS) and 1x penicillin/streptomycin and were maintained in a humidified incubator at 37°C with 5% CO_2_. Transfection was performed with Lipofectamine 3000 (Invitrogen) according to manufacturer’s instructions, 0.3-0.6 μg DNA/coverslip were used and cells were analysed 18-48 h post-transfection.

### Microfluidics chambers

Microfluidic chambers (MFC) were made with Sylgard 184 silicone elastomer kit (Dow Corning) using epoxy resin moulds previously designed in the laboratory (Restani et al., 2012). Once the MFCs were baked, reservoirs were cut and the MFCs were mounted onto glass bottom dishes (HBST-5040, WillCo well), pre-coated with 20 μg/ml poly-D-Lysine. MFCs were then blocked with 0.8% ES grade BSA (Sigma) overnight, poly-ornithine (>3 h) and finally laminin (overnight), before plating motor neurons. MFCs have 500 μm long grooves that separate the somatic from the axonal compartment.

### Constructs

mCherry-FUS^513x^ (ΔNLS) was kindly gifted by D. Dormann, pcDNA6-Flag-FUS was gifted by M.-D. Ruepp. pcDNA6-Flag-FUS^P525L^ was generated in the lab by PCR mutagenesis. pBABE-puro SV40 LT was a gift from T. Roberts (Addgene plasmid # 13970; http://n2t.net/addgene:13970; RRID:Addgene_13970).

### Antibodies

The following antibodies were used: anti-FUS N-term (WB 1:5000, NB100-565, Novus Biologicals), anti-FUS C-term (IF 1:300, WB 1:5000, NB100-562, Novus Biologicals), anti-Δ14 (IF 1:300, WB 1:1000 (Devoy et al., 2017)), β3-tubulin (1:1000, cat. no. 801202, Biolegend; 1:500, cat. no. 302 306, SySy) anti-GFP (IF 1:1000, GFP1011, Aves labs), anti-FMRP (IF 1:300, WB 1:1000, ab17722, Abcam), anti-SMN1 (IF 1:300, WB 1:1000, cat. no. 610646, BD), anti-Flag M1 (IF 1:500, cat. no. F3040, Sigma), anti-G3BP1 (1:200, cat. no. 611126, BD), anti-RPL26 (IF 1:800, WB 1:2000, ab59567, Abcam), anti-RPS6 (WB 1:1000, Cell Signalling), anti-LAMP1 (IF 1:300, ab25245, Abcam), anti-GAPDH (WB 1:5000, mab374, Millipore), anti-HA (WB 1:3000, IHC: 1:100 cat. no. H6908, Sigma-Aldrich), anti-ChAT (IHC 1:100, Chemicon). AlexaFluor conjugated secondaries were from Invitrogen (1:1000) or Jackson ImmunoResearch (1:500).

### Immunofluorescence and image analysis

Cells were fixed in a PFA solution (4% PFA, 4% sucrose in PBS) for 15 min at room temperature (RT). Samples were then permeabilised and blocked in a solution containing 10% HRS, 0.5% BSA, 0.2% Triton X-100 in PBS for 15 min. Primary antibodies were diluted in blocking solution (10% HRS, 0.5% BSA in PBS) and incubated for 1 h at RT. Secondary antibodies were diluted in blocking solution (10% HRS, 0.5% BSA in PBS) and incubated for 1 h at RT.

Coverslips were mounted using Mowiol or FluoromountG (Thermo Fisher); MFCs with Ibidi mounting media. Imaging was carried out using a Zeiss LSM 780 inverted confocal microscope with a 40x oil-immersion lens with 1.3 numerical aperture, or with a Zeiss LSM 710 inverted confocal microscope with a 63x oil-immersion lens with 1.4 numerical aperture. Images were digitally captured using ZEN 2010 software and analysed using Fiji (ImageJ). Digital deconvolution was performed using ImageJ plug-ins. ‘Diffraction PSF 3D’ was used to generate a theoretical point spread function for each wavelength, and ‘Parallel spectral deconvolution 2D’ was used for the generation of the deconvolved image. SynPAnal (Danielson and Lee, 2014) was used to quantify axonal puncta size.

### Immunohistochemistry

For immunohistochemical analysis three months old mice were perfused with saline followed by 4% PFA solution. Spinal cords were dissected, incubated in 20% (wt/vol) sucrose, embedded in Tissue-Tek OCT compound (Sakura Finetek, 4583) and sectioned with an OTF Cryostat (Bright Instruments). Slices were mounted on microscope slides and sections were encircled with a hydrophobic barrier pen (Dako, S2002), permeabilised by three 10 min washes with 0.3% Triton X-100 in PBS, and blocked for 1 h in 10% BSA and 0.3% Triton X-100 in PBS. Samples were then probed with primary antibodies overnight, the samples were then washed three times prior to incubation with secondary antibodies for 1 h. Slides were then washed, mounting media added and samples were covered with 22 × 50 mm cover glass.

### Cellular translation assay

L-azidohomoalanine (AHA) labelling assays were carried out as previously described (Moens et al., 2019). Briefly, neurons were incubated in a neuronal methionine-free media consisting of: methionine and cysteine free DMEM supplemented with 0.26 mM L-cysteine, 0.23 mM sodium pyruvate, 10 mM HEPES pH 7.4, 0.067 mM L-proline, 0.674 μM zinc sulphate, 5nM B12, 1x Glutamax, 1x B27, 1x penicillin/streptomycin, 1 ng/mL BDNF, 0.1 ng/mL GDNF, 10 ng/mL CNTF for 30 min prior to the addition of 2 mM AHA or vehicle control for 30 min. Anisomycin (40 μM) was pre-incubated for 20 min and co-incubated with AHA. Neurons were then fixed, permeabilised and AHA was labelled by click chemistry using Click-iT Cell Reaction Buffer Kit with an AlexaFluor-555 alkyne (1μM) following the manufacturer’s instructions.

### Proximity Ligation Assays

Proximity ligation assay was performed with Duolink® In Situ Orange PLA reagents according to the manufacturer’s protocol (Sigma Aldrich).

### Western blotting and co-immunoprecipitation

Motor neuron cultures were lysed in RIPA buffer (50 mM Tris–HCl pH 7.5, 150 mM NaCl, 1% NP-40, 0.5% sodium deoxycholate, 0.1% SDS, 1 mM EDTA, 1 mM EGTA, Halt™ phosphatase and protease inhibitor cocktail (Thermo Fisher)), incubated on a rotating wheel at 4 °C for 1 h, and then nuclei and cellular debris were spun down at 20,000xg for 10 min. Supernatants were collected, Laemmli buffer was added and samples were denatured at 98°C for 5 min.

For co-immunoprecipitation (co-IP) assays E12.5-14.5 brains were used. Samples were homogenised in lysis buffer (20 mM HEPES pH 7.4, 150 mM NaCl, 10% glycerol, Halt™ phosphatase and protease inhibitor cocktail), incubated on a rotating wheel at 4 °C for 1 hour, nuclei and cellular debris were spun down at 20,000xg for 20 min. Supernatants were collected, 1mg of protein lysate was used per co-IP and protein of interest immunoprecipitated with 2 μg of antibody or appropriate IgG control overnight. Protein A-Sepharose beads (Sigma-Aldrich) were used to purify the antibody/protein complex, precipitates were washed three times prior to being eluted in Laemmli buffer.

Samples were separated on precast 4–15% Mini-PROTEAN® TGX Stain-Free™ Protein gels (Bio-Rad) and transferred onto a polyvinylidene difluoride (PVDF) membrane using a semi-dry Trans-Blot Turbo system (Bio-Rad), or NuPAGE™ 4-12% Bis-Tris Protein gels were used, and proteins blotted onto nitrocellulose membrane using a Novex system (GE Healthcare). Western blots were developed with Classico substrate (Millipore), and detected with a ChemiDoc imaging system (Bio-Rad). Densitometric quantification of bands was carried out using the software Image Lab (Bio-Rad).

### Polysome profiling

Cytoplasmic lysates from frozen E17.5 bains were prepared as described previously (Bernabò et al., 2017). Tissue was pulverised in a mortar under liquid nitrogen. The tissue powder was dissolved in 10 mM Tris-HCl pH 7.5, 10 mM NaCl, 10 mM MgCl_2_, 1% Triton X-100, 1% sodium deoxycholate, 0.4 U/ml RiboLock RNase Inhibitor (Thermo Scientific), 1 mM dithiothreitol, 0.2 mg/ml cycloheximide, 5 U/ml DNase I (Thermo Scientific). Following a first centrifugation step for 1 min at 14000g at 4°C to remove tissue debris, the supernatant was centrifuged for 5 min at 14000 to pellet nuclei and mitochondria. Cleared supernatants were then loaded on a linear 15%–50% sucrose gradient in 10 mM Tris-HCl pH, 100 mM NaCl, 10 mM MgCl_2_ and ultracentrifuged in a SW41Ti rotor (Beckman) for 1 h and 40 min at 40,000 rpm at 4°C in a Beckman Optima LE-80K Ultracentrifuge. After ultracentrifugation, gradients were fractionated in 1 mL volume fractions with continuous monitoring absorbance at 254 nm using an ISCO UA-6 UV detector. Proteins were extracted from each sucrose fraction of the profile using the methanol/chloroform protocol and solubilised in a sample buffer.

### Cloning and purification of MBP-FUS and mutants

MBP-FUS was a gift from Nicolas Fawzi (Addgene plasmid # 98651; http://n2t.net/addgene:98651; RRID:Addgene_98651). The P525L FUS point mutation and the Δ14 mutant version of FUS were generated via site-directed mutagenesis using MBP-FUS. The amino acid sequence at the C-terminus of the Δ14 mutant is “KAPKPDGPGGGPGGSHMGVSTDRIAGRGRIN*”.

MBP-FUS and mutants were expressed in *E. coli* BL21 DE3 cells with rare codons for R,I,P and L using chloramphenicol and kanamycin for selection. Following cell lysis by sonication, the protein was purified by nickel-affinity chromatography. The lysis buffer used contained 20 mM sodium phosphate, 500 mM NaCl, 5 mM β-mercaptoethanol and 20 mM imidazole, pH 7.4. One Complete Protease Inhibitor tablet (Sigma-Aldrich) was added to the lysate from 2 litres of growth. The column was washed with the same buffer supplemented with 40 mM imidazole. Protein was eluted with lysis buffer with 400 mM imidazole. The protein was then further purified using gel filtration chromatography with a buffer containing 50 mM Tris-HCl pH 7.6, 150 mM NaCl, 5 mM β-mercaptoethanol and 1 mM EDTA.

### Protein expression and purification of FMRP-Cterm

The low-complexity disordered region of human FMRP_445-632_ (referred to as FMRP_LCD_) was expressed and purified as previously described (Tsang et al., 2019). Briefly, His-SUMO-FMRP was transformed into *E. coli* BL21-CodonPlus(DE3) RIL cells and grown at 37°C in LB. Protein expression was induced with 0.5 mM IPTG at an OD_600nm_ ∼0.6 and grown overnight at 25°C. Cells were harvested and lysed in lysis buffer containing 6 M guanidinium chloride (GdnHCl), 50 mM Tris-HCl pH 8.0, 500 mM NaCl, 20 mM imidazole and 2 β-mercaptoethanol. Harvested cells were sonicated for 4.5 mins (2 s on, 1 s off), and centrifuged. The supernatant of the lysate was then purified by nickel-affinity chromatography equilibrated with the lysis buffer. The column was washed in lysis buffer without 6 M GdnHCl, then eluted in buffer containing 50 mM Tris-HCl pH 8.0, 500 mM NaCl, 300 mM imidazole and 2 mM mM β-mercaptoethanol. The His-SUMO tag was cleaved with ULP protease while dialyzed against cleavage buffer (50 mM Tris-HCl, 150 mM NaCl, 20 mM imidazole, 2mM mM β-mercaptoethanol at pH 7.4) overnight at 4°C. FMRP was separated from the His-SUMO tag by nickel-affinity chromatography following the same steps described above. The fractions containing FMRP were collected and successful separation of FMRP from the His-SUMO tag was verified with SDS-PAGE gel. FMRP was concentrated and further purified using gel filtration chromatography with a buffer containing 4 M GdnHCl, 50 mM Tris-HCl pH 8.0, 500 mM NaCl, and 2 mM β-mercaptoethanol.

### Fluorescence protein labeling

An AlexaFluor-647 fluorescent dye was added to the only cysteine (C584) in FMRP_445-632_ via a maleimide linkage following manufacturer’s instruction with slight modifications. First, FMRP was dialyzed into a buffer containing 50 mM Tris-HCl pH 7.5, 100 mM NaCl and 4 M GdnHCl. To ensure that any residual reducing agents were removed, the protein was desalted using a Hi-Trap desalting column (GE Healthcare). After desalting, the protein sample was immediately reacted with 5x AlexaFluor-647 (ThermoFisher) maleimide dye. The reaction was incubated overnight at 4°C and quenched with an excess of reducing agent (DTT) the following day. To remove any unreacted dye, the protein was passed through a Hi-Trap desalting column (GE Healthcare) and an S75 gel filtration column equilibrated in buffer containing 50 mM Tris-HCl pH 7.5, 100 mM NaCl, 4 M GdnHCl and 2mM DTT. Successful dye separation was confirmed by running the protein sample on an SDS-PAGE gel and then visualizing any remaining free dye with a fluorescence reader ChemiDoc MP System (BioRad). Labeled proteins were either frozen or dialyzed into specific assay buffers.

### RNA Preparation

Sc1 RNA (GCUGCGGUGUGGAAGGAGUGGUCGGGUUGCGCAGCG) was purchased from Sigma-Aldrich as lyophilized samples. 100 μM stocks were reconstituted in water and stored at −20°C. Working stocks were diluted into specific assay buffers.

### Turbidity measurements

For turbidity measurements, OD_600nm_ of MBP-FUS was obtained using a SpectraMax i3x Multi-Mode Plate Reader (Molecular Devices) at 25°C. The samples were prepared by mixing varying concentrations of MBP-FUS with 0.5 μM TEV protease in a buffer containing 25 mM Tris-HCl pH 7.4, 150 mM KCl and 2 mM DTT. Samples were equilibrated for 5 min before reading the turbidity. Turbidity was measured at intervals of 35 s for a total of 20 min. The change in turbidity was calculated from the slope (Δ absorbance/min) from 0 to 5 min. Apparent C_sats_ are calculated as previously described (Wang et al., 2018).

### In vitro co-LLPS assays

*In vitro* phase separation assays of low complexity regions of FUS and FMRP were performed using ^FITC^FUS^LCD^ (50μM) and ^Alexa647^FMRP^LCD^ (50μM) in the presence of 1 μM sc1 RNA in a buffer containing 25 mM sodium phosphate pH 7.4, 50 mM KCl, 2 mM DTT. For partitioning assays using full length FUS proteins (FUS^WT^, FUS^P525L^ and FUS^Δ14^), MBP-FUS (10 μM) phase separation was induced by TEV protease (0.5 μM) cleavage before the addition of ^Alexa647^FMRP^LCD^ (1 μM) and sc1 RNA (0.5 μM) in a buffer containing 25 mM sodium phosphate pH 7.4, 150 mM KCl, 2 mM DTT.

### Fluorescence microscopy of phase separated samples

Fluorescence images of phase separated droplets were imaged on a confocal Leica DMi8 microscope equipped with a Hamamatsu C9100-13 EM-CCD camera with a 63x objective. AlexaFluor-647 fluorescence was detected using a 637 nm laser and FITC fluorescence was detected using a 491 nm laser. In experiments with MBP-FUS and MBP-FUS mutants, samples were incubated with TEV protease for 10 min before imaging. All phase-separated droplets were imaged on a 96 glass well plate (Eppendorf). Two-or three-fold concentrated protein or RNA samples were prepared to account for the dilution in mixing with other components to achieve desired final concentrations. Note that no molecular crowding reagents were used. Images represent droplets settled to the bottom of the plate. Images were processed using Volocity (Perkin Elmer) and ImageJ.

### In vitro partitioning assay

To determine the partitioning of FMRP, images of droplets with the addition of 5% AlexaFluor-647-FMRP_LCD_ were acquired as described above and analyzed with ImageJ. An image of the buffer in the absence of any protein was used to subtract any background artefacts. In ImageJ, masks were defined using the Otsu threshold method while applying several criteria to the particle picking algorithm: droplets are required to have a radius greater than 1 μm, and with the circularity of 0.5-1.0. The intensity of the bulk background solution is defined as the mean intensity within a circular region of interest with a diameter of 5 μm that does not contain any phase separated droplets. Fluorescence enrichment ratios were calculated from the ratio of the mean fluorescence intensity (inside droplet) / mean fluorescence intensity (background outside of droplet). Droplets were randomly imaged and measurement represent three independent experiments.

### In vitro translation assay

In vitro translation rates represent the increase in luminescence as a function of time using a standard rabbit reticulocyte lysate system (Promega) with luciferase mRNA (Promega). Manufacturer’s instructions were followed with a few modifications. Briefly, each reaction (30 μL) contains 12.6 μL of rabbit reticulocyte lysate, 0.5 μL of luciferase mRNA (1 mg/mL), 0.3 μL of amino acid mixture minus leucine (1 mM), 0.3 μL of amino acid mixture minus methionine (1 mM), 2 uL of TEV protease (15 μM) and 14.3 μL of 5 μM protein (MBP-FUS/FMRP) or buffer (25 mM sodium phosphate pH 7.4, 50 mM KCl and 2 mM DTT). First, the reaction was incubated for 10 min, then end-point luminescence measurements were carried out in intervals of 10 min up to 50 min. Each end-point luminescence measurement contained 75 μL of luciferase substrate mixed with 2.5 μL of unpurified translation mixture measured in a white opaque 96 well plate (Corning 3990). A SpectraMax i3x Multi-Mode Plate Reader (Molecular Devices) at 25 °C was used to detect the luminescence. The translation rates represent the line of best fit from the end-point luminescence readings as a function of time.

### RiboTag

The RiboTag method was performed as described previously (Shigeoka et al., 2016) with modifications. Briefly, E17.5 spinal cords were homogenised using Tissue Ruptor (Qiagen) in a buffer (50 mM Tris-HCl pH 7.4, 100 mM KCl, 12 mM MgCl_2,_ 1% NP-40) supplemented with 0.1 mg/ml cycloheximide, 1 mg/ml heparin, SuperaseIn RNAse (Thermo Fisher). Lysates were cleared by centrifugation at 10000xg for 10 min and 5% of the lysate was saved as input. To reduce nonspecific binding protein G magnetic beads (DynaBeads, Thermo Fisher) were added to the lysate and incubated for 2 hours at 4°C. Next, 5 ul of anti-HA antibody (Sigma) was added to the precleared lysate and incubated for 2 hours at 4 °C. Later 100 μl beads slurry with 2 μl of SuperaseIn were added to the lysate followed by 2h incubation in the cold room. After precipitation, beads were washed 5 times in the wash buffer (300 mM KCl, 1% NP-40, 50 mM Tris-HCl pH 7,4, 12 mM MgCl_2_, 0.1 mg/ml cycloheximide). Beads were eluted in Qiazol (Qiagen) and RNAs were isolated using RNeasy Micro Kit (Qiagen). 10% of the beads were used for WB and eluted in Laemmli sample buffer (Thermo Fisher) supplemented with 100 mM DTT. The quality control of RNA was performed using TapeStation (Agilent).

### qPCR analysis of RiboTag samples

SuperScript IV VILO (Thermofisher) was used to reverse transcribe RNA from input and IP samples (concentrations adjusted). cDNA was used for qPCR analysis, expression of GAPDH was used as housekeeping control. Primers used for qPCR analysis: Chat forward: GCGTAACAGCCCAGGAGAG, Chat reverse: TTGTACAGGCATCTTTGGGG, Gapdh forward: CAAGCTCATTTCCTGGTATGA, Gapdh reverse: CTCTTGCTCAGTGTCCTTGCT, HA-Rpl22 forward: GTGCCTTTCTCCAAAAGGTATTT, HA-Rpl22 reverse: GTCATATGGATAGGATCCTGCATA. Pmp22 forward: GCCGTCCAACACTGCTACTC, Pmp22 reverse: GAGCTGGCAGAAGAACAGGA.

### RNA Sequencing and analysis of RiboTag samples

Libraries were prepared using NEBNext mRNA Ultra II in UCL Genomics facility and sequenced (75bp single-end) to an average depth of 18 million reads. Each sample was aligned to the *Mus musculus* (house mouse) genome assembly GRCm38 (mm10) with STAR (v2.4.2a) (Dobin et al., 2013). Reads were coordinate sorted and marked for PCR duplicates using Novosort (1.03.09). Gene expression was quantified using HTSeq using the Ensembl mm10 (v82) mouse transcript reference (Mudge and Harrow, 2015). Differential gene expression was calculated using DESeq2 (Love et al., 2014) comparing the IP samples between the *Fus*^-/-^ (n=4) and *Fus*^Δ14/Δ14^ (n=5) with the same number of their respective littermate controls, in two separate analyses. The significance level was set at a false discovery rate adjusted p-value of 10%. To compare the two analyses, each nominal p value was converted into a Z-score and given the sign of the log_2_ fold change. We defined condition-specific genes as having adjusted p value of < 0.1 in one condition and > 0.1 in the other. A list of FMRP target genes was obtained from a published HITS-CLIP experiment (Darnell et al., 2011). Entrez IDs were converted to Ensembl IDs using g:Convert from the g:Profiler suite of tools (Reimand et al., 2007). A list of FUS target peaks were obtained from a published iCLIP experiment (Rogelj et al., 2012). Peaks were annotated using annotatr (Cavalcante and Sartor, 2017) using gene models included in the TxDb.Mmusculus.UCSC.mm9.knownGene and org.Mm.eg.db R packages (Carlson et al., 2015, 2019). FUS targets were filtered to only use peaks overlapping coding and UTR regions.

### Statistical analysis

Unless otherwise stated data were obtained using cells from at least three independent preparations, which are visualised in different shades of grey in the graphs. The numbers of cells studied are given in the figure legends. GraphPad Prism or R were used for statistical analysis. Normality of data distribution was tested using D’Agostino and Pearson normality test. One-way ANOVA followed by Dunnet’s post hoc test was used for normally distributed data and multiple comparisons, Kruskal-Wallis followed by Dunn’s post hoc was used for not normally distributed data and multiple comparisons. Friedman’s test, followed by Dunn’s post hoc test was used to compare normally distributed paired samples. Individual differences were assessed using individual Student’s t tests. Data are shown as mean ± SEM. Kolmogorov-Smirnov test was used in cumulative frequency analysis to test differences between targets and non-targets of FUS and FMRP.

## Author contributions

Conceptualization, P.F.; Investigation, N.B, A.M.U, M.G.G., B.T., F.M., M.P., O.G.W, G.V.; Data Curation, J.H, S.J.; Formal Analysis, A.M.U., N.B.; Writing – Original Draft, N.B., P.F., A.M.U.; Writing – Review & Editing, N.B., A.M.U., M.G.G., B.T., F.M., P.A.C., J.H., E.M.C.F., A.R., G.V., J.D.F.-K., G.S., P.F.; Resources, C.B., R.D.L.F., B.T., P.A.C., M.N., A.D.; Funding Acquisition, P.F., J.D.F.-K., G.V.; Supervision, P.F., J.D.F.-K., G.S.

## Acknowledgements

We thank Dorothee Dorman and Marc-David Ruepp for valuable discussion. Emily Spaulding and Robert Burgess for helpful discussion and technical advice on the RiboTag method. James Sleigh for critical reading of this manuscript; and all members of the Fratta and Schiavo labs for useful discussion.

PF is funded by an MRC/MNDA Lady Edith Wolfson Fellowship and by the NIHR-UCLH Biomedical Research Centre. This work was funded by UK Medical Research Council (MR/M008606/1 and MR/S006508/1 to PF; MR/S022708/1 to EMCF and PF); Motor Neurone Disease Association (885-792 to PF); Rosetrees Trust (to PF and EMCF); Wellcome Trust Senior Investigator Award (107116/Z/15/Z) (GS), the UK Dementia Research Institute Foundation award (UKDRI-1005) (GS), the Canadian Institutes of Health Research Foundation Grant #148375 (JDF-K) and AriSLA Foundation (AxRibALS) (GV).

## SUPPLEMENTARY FIGURES

**SFig. 1.**
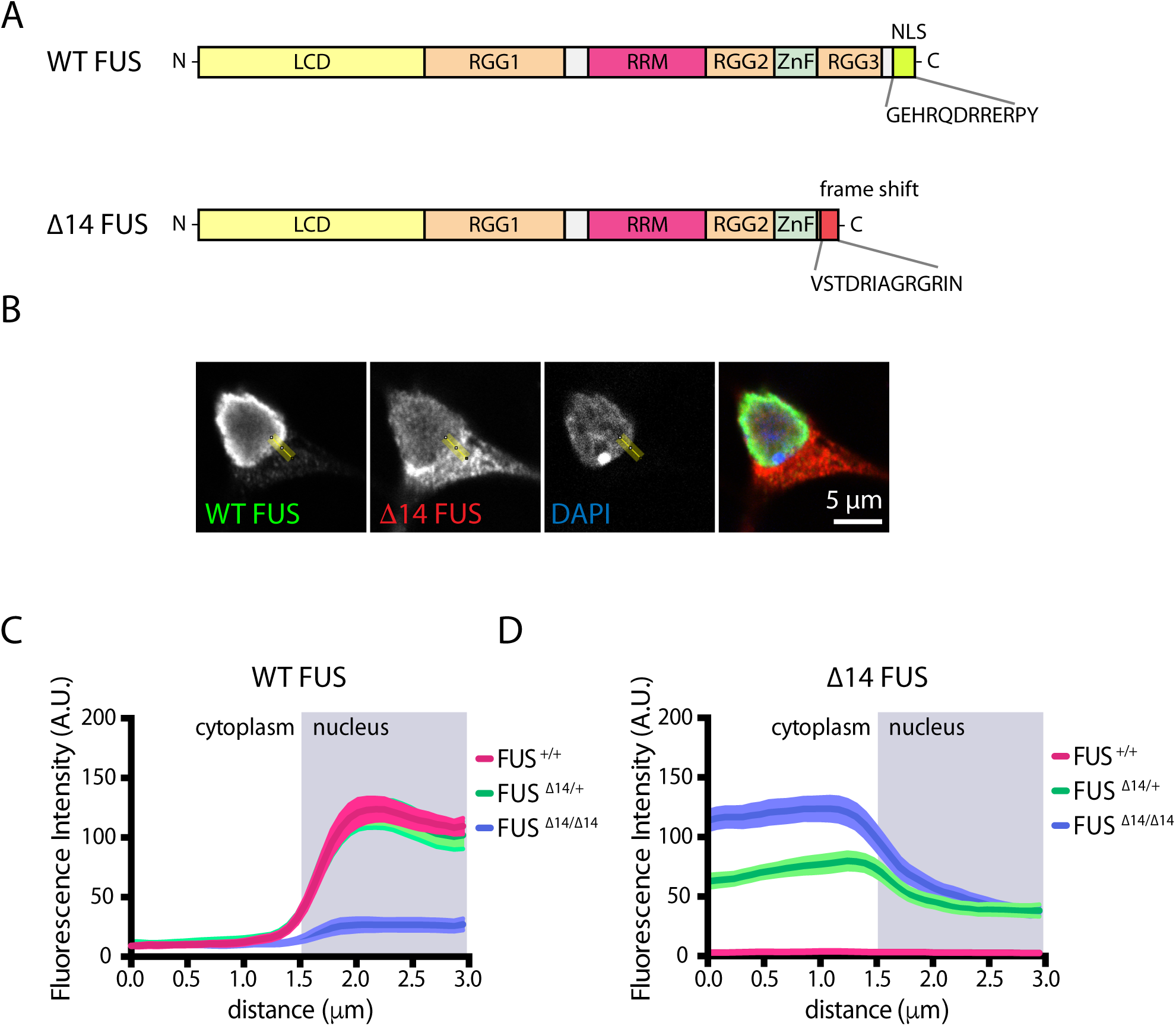
FUS localisation in Δ14 FUS expressing primary MNs. **(A)** Schematic representation of wild type and Δ14 FUS domains. **(B, C, D)** Analysis of the localisation of wild type and Δ14 mutant FUS. **(B)** Example of a *Fus^Δ14/+^* MN with a 3 μm line (ROI) crossing the nucleus and cytoplasm. **(C, D)** Profile of wild type (C) or Δ14 (D) FUS fluorescence intensity detected in the ROI in *Fus*^+/+^ (pink), *Fus^Δ14/+^* (green) and *Fus^Δ14/Δ14^* (blue) MNs (n=5, MNs=40).

**SFig. 2.**
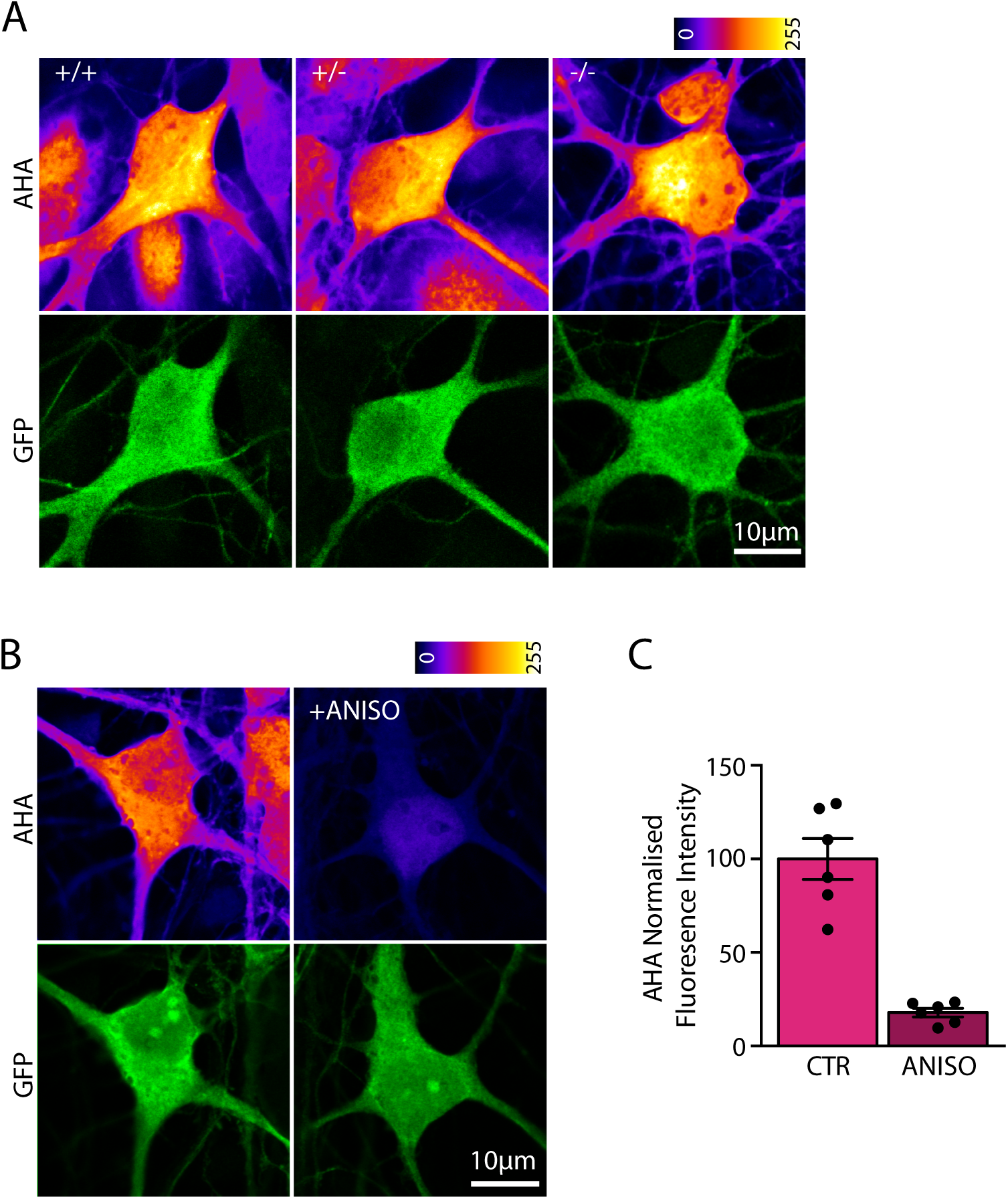
Translation is unaffected in FUS knock-out MNs. **(A)** Metabolic labelling using the methionine analog L-azidohomoalanine (AHA, 2 mM, 30 min) and click chemistry in primary *Fus*^+/+^, *Fus*^+/-^ and *Fus*^-/-^ motor neurons. AHA labelling is visualised using the LUT fire (top panels), motor neurons are identified by the GFP expression under the HB9 promoter (bottom panels). **(B,C)** AHA signal (LUT fire, top panels) is blocked by pre-incubation with the translation inhibitor anisomycin (20 min, 40 μM, right panel). **(C)** Quantification of the effect (n=1, MNs=6).

**SFig. 3.**
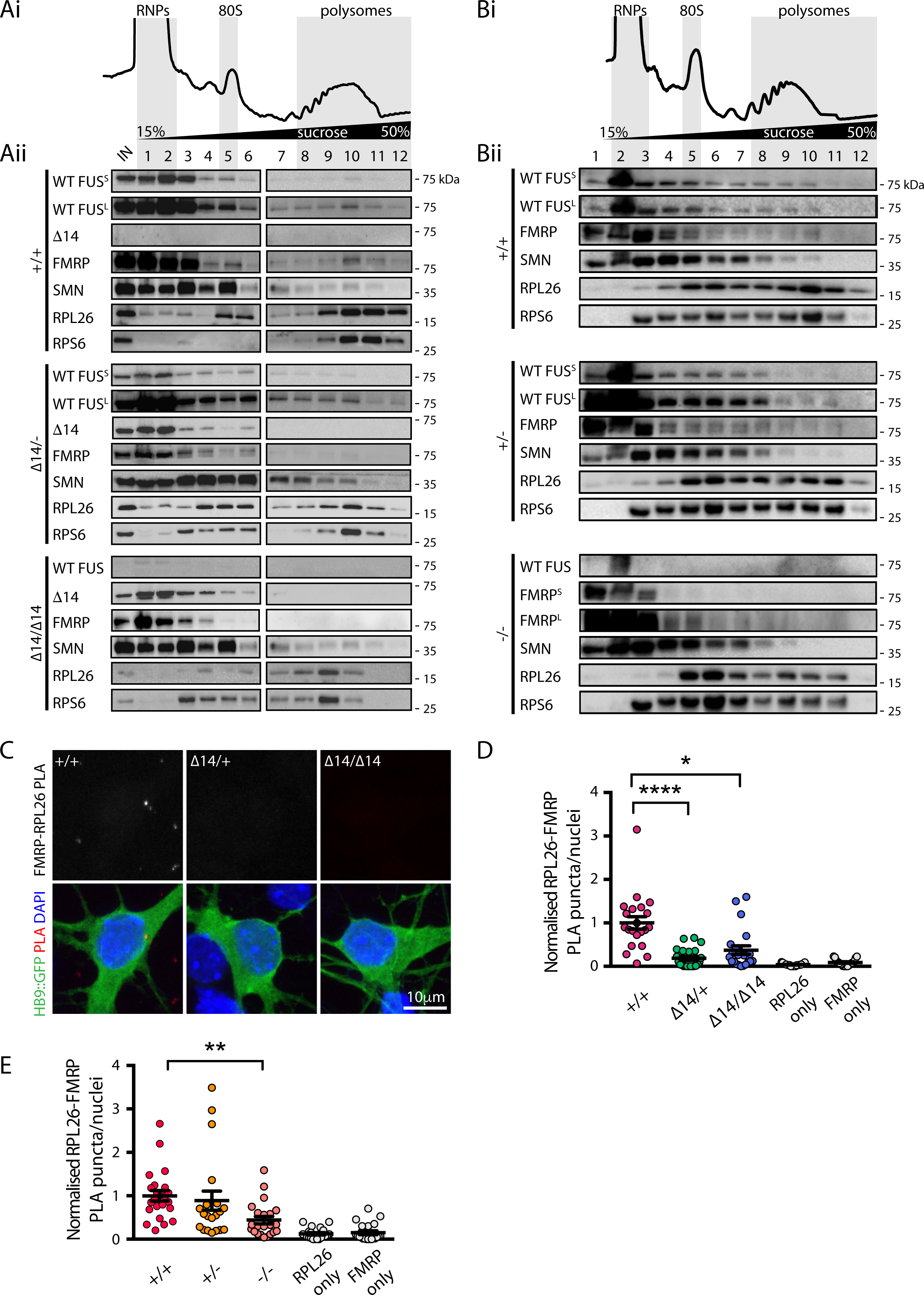
Polysome association of mutant FUS and FMRP in *Fus^Δ14/Δ14^* and *Fus*^-/-^. **(A)** Representative polysome (i) and co-sedimentation (ii) profiles of *Fus^+/+^*, *Fus^Δ14/+^* and *Fus^Δ14/Δ14^* E17.5 mouse brains. **(Ai)** From left to right: free cytosolic fraction (RNPs), monosomes (80S) and polysomal fractions. **(Aii)** Western blotting of the RBPs of interest, RPL26 and RPS6 were used as co-sedimentation controls (S=short exposure, L=long exposure, n=2). **(B)** Representative polysome (i) and co-sedimentation (ii) profiles of *Fus*^+/+^, *Fus*^+/-^ and *Fus*^-/-^ E17.5 mouse brains. **(Bii)** Western blotting of the RBPs of interest, RPL26 and RPS6 were used as co-sedimentation controls (S=short exposure, L=long exposure, n=2). **(C)** Proximity ligation assay (PLA) detecting proximity between FMRP and RPL26 (top panel) in *Fus^+/+^*, *Fus^Δ14/+^* and *Fus^Δ14/Δ14^* primary motor neurons identified by GFP expression (HB9::GFP). **(D)** Quantification of FMRP-RPL26 PLA puncta density in *Fus^+/+^*, *Fus^Δ14/+^* and *Fus^Δ14/Δ14^* MNs normalised to wild type (n=4, 21-22 images; *p=0.0431, ****p<0.0001, Kruskal-Wallis followed by Dunn’s post hoc test). **(E)** Quantification of FMRP-RPL26 PLA puncta density in *Fus^+/+^*, *Fus^+/-^* and *Fus^-/-^* MNs normalised to wild type (n=4, 20-23 images; **p<0.001 Kruskal-Wallis followed by Dunn’s post hoc test).

**SFig. 4.**
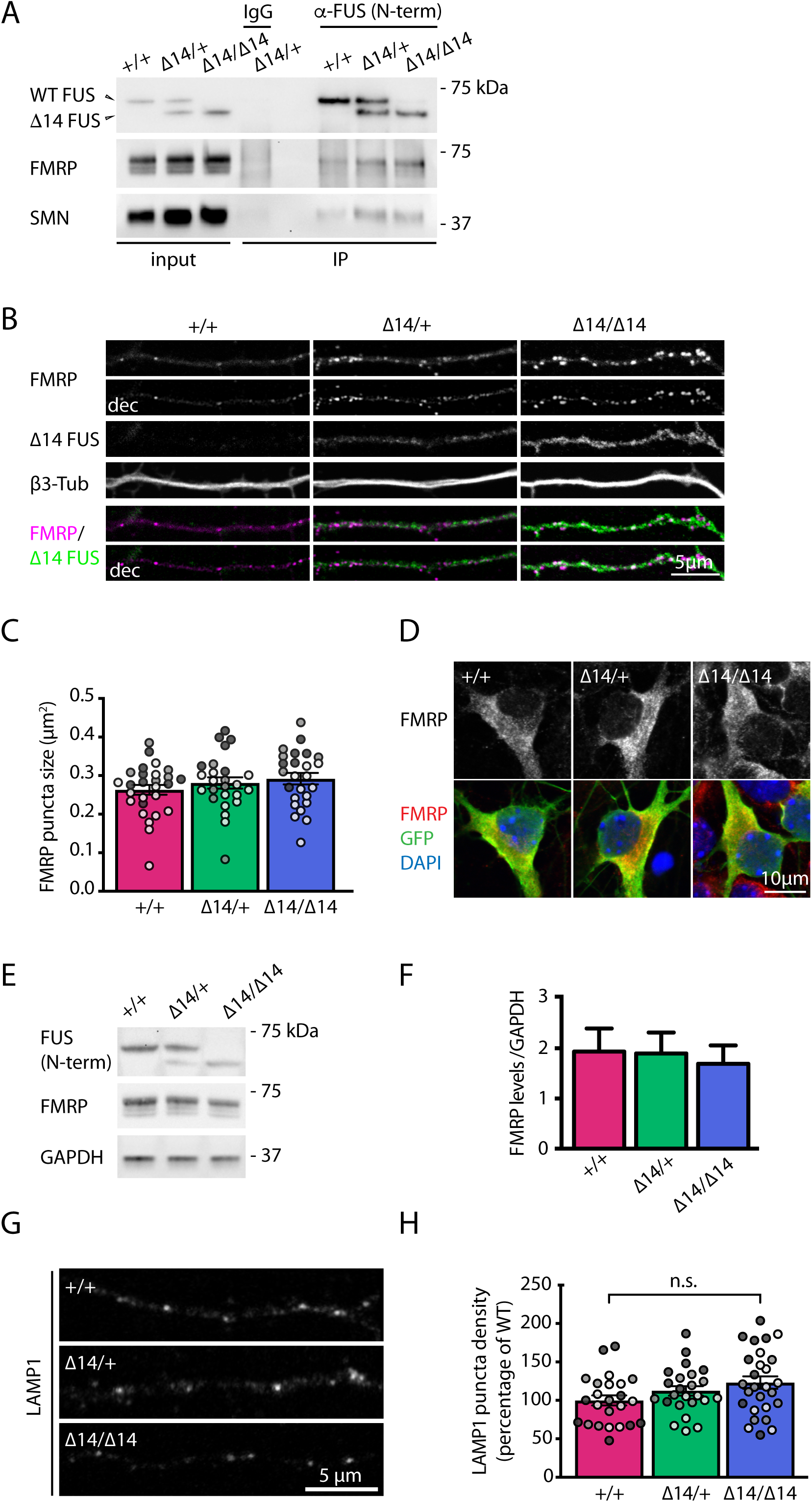
Axonal FMRP puncta density is increased in Δ14 FUS MNs despite no changes in total protein expression. **(A)** Co-immunoprecipitation assay. Wild type or Δ14 FUS are immuno-purified from *Fus*^+/+^, *Fus^Δ14/^*^+^ and *Fus^Δ14/Δ14^* embryonic E13.5 brains with a N-terminal FUS antibody. FMRP and SMN both co-immunoprecipitate with WT and mutant FUS. Rabbit IgGs are used as a control, input on the left shows similar expression. **(B)** Representative images of FMRP (magenta) and Δ14 FUS (green) labelling in *Fus^+/+^*, *Fus*^Δ14/+^ and *Fus^Δ14/Δ14^* MNs axons grown in MFCs (DIV 8). Both original and deconvolved FMRP images are shown. β3-tubulin is used to identify axons. **(C)** FMRP puncta size analysis in *Fus*^+/+^, *Fus*^Δ14/+^ and *Fus^Δ14/Δ14^* MNs axons (n=3, axons=38-41). **(D)** Representative images of somatic FMRP labelling in primary *Fus*^+/+^, *Fus^Δ14/+^* and *Fus^Δ14/Δ14^* MNs identified by HB9 driven GFP expression. **(E)** Representative western blot showing expression levels of FMRP in *Fus*^+/+^, *Fus*^Δ14/+^ and *Fus^Δ14/Δ14^* primary MN cultures. Wild type and Δ14 FUS are probed with a N-terminal FUS antibody. Wild type FUS bands have a higher molecular weight (∼70 kDa) compared to Δ14 FUS. GAPDH is used as a loading control. **(F)** Quantification of FMRP expression levels, normalised to GAPDH. (n=3) **(G)** Axonal LAMP1 staining in *Fus*^+/+^, *Fus*^Δ14/+^ and *Fus^Δ14/Δ14^* MNs grown in MFCs. **(H)** Quantification of LAMP1 puncta density as shown in (G) and normalised to *Fus*^+/+^ density (n=3, MNs=24-27). Independent experiments are visualised in different shades of grey in the graphs.

**SFig.5.**
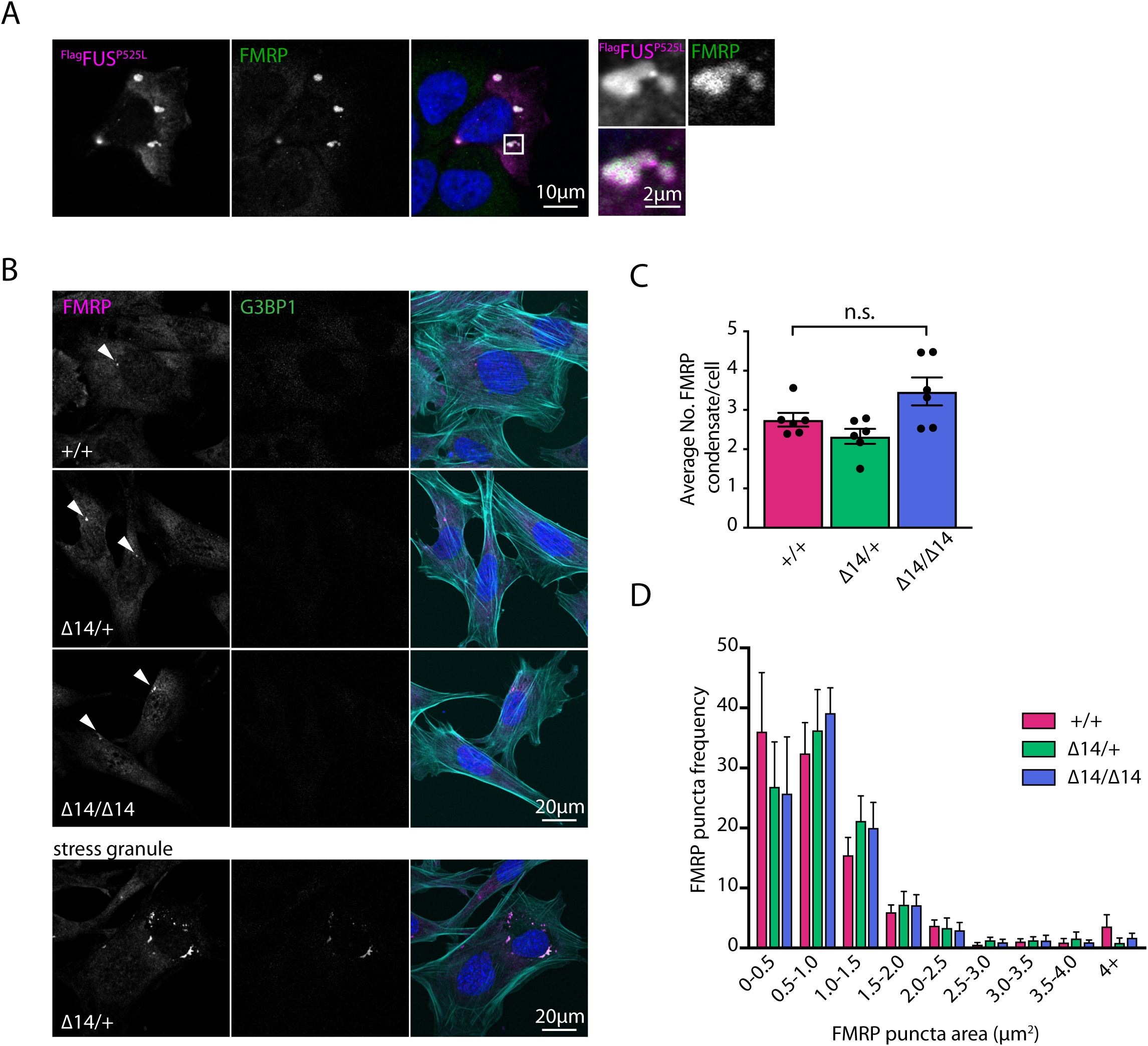
Mutant FUS expression drives the formation of FMRP condensates. **(A)** HeLa cells overexpressing ^Flag^FUS^P525L^. Endogenous FMRP (middle panel, green) is recruited to ^Flag^FUS^P525L^ condensates. On the right, zoomed images of the condensates in the white box (merged image). **(B)** Representative images of *Fus^+/+^*, *Fus*^Δ14/+^ and *Fus^Δ14/Δ14^* MEFs with FMRP puncta (>0.5μm^2^, white arrowheads). FMRP puncta are mostly negative for the stress granule marker G3BP1 (middle panel). Phalloidin (cyan) is used to identify cells and DAPI (blue) to label nuclei. Bottom panels show an example of a *Fus^Δ14/+^* cell with FMRP and G3BP1 positive granules (stress granules). **(C)** Quantification of the number of FMRP condensates per cell in *Fus*^+/+^, *Fus*^Δ14/+^ and *Fus*^Δ14/Δ14^ MEFs (n=6). **(D)** Analysis of FMRP puncta size in *Fus*^+/+^, *Fus*^Δ14/+^ and *Fus^Δ14/Δ14^* MEFs (n=6).

**SFig.6.**
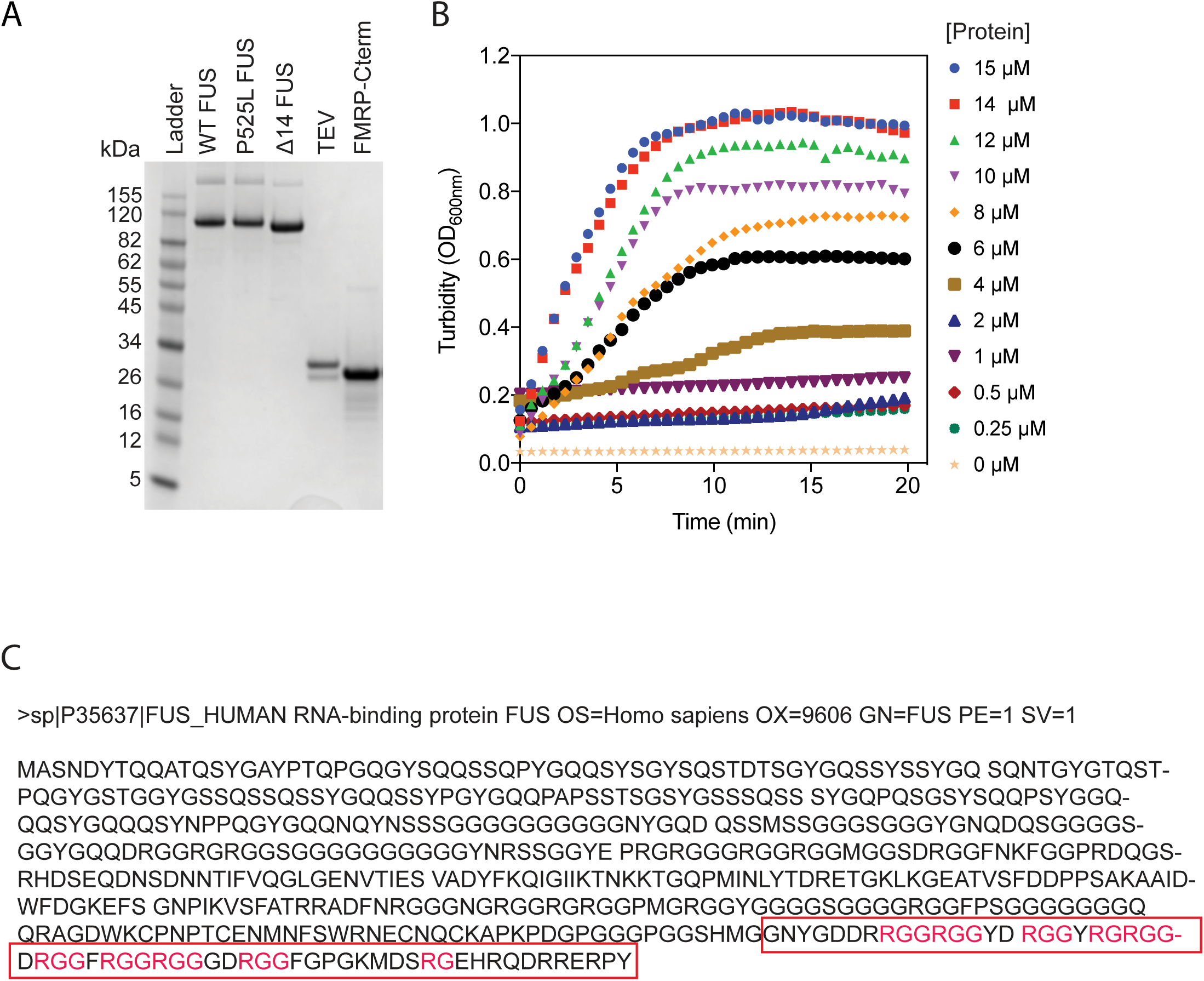
Characterisation of recombinant proteins used for *in vitro* assays and example of turbidity measures. **(A)** Coomassie stained SDS-PAGE gel of *in vitro* purified proteins used in the study. FUS proteins are tagged with an N-terminal MBP. **(B)** Example of a turbidity progress curve for FUS phase separation. Change in turbidity (OD 600_nm_) of different FUS concentrations was measured as a function of time. Turbidity was induced by addition of 500 nM of TEV. The rate of change in turbidity was determined as the slope of the initial period of turbidity change (0 to 6 minutes) for each protein concentration. **(C)** Δ14 FUS deletion (deleted region boxed in red) results in loss of ∼1/2 of total RG/RGG motifs (bolded in red) in the C-terminal region.

**SFig. 7.**
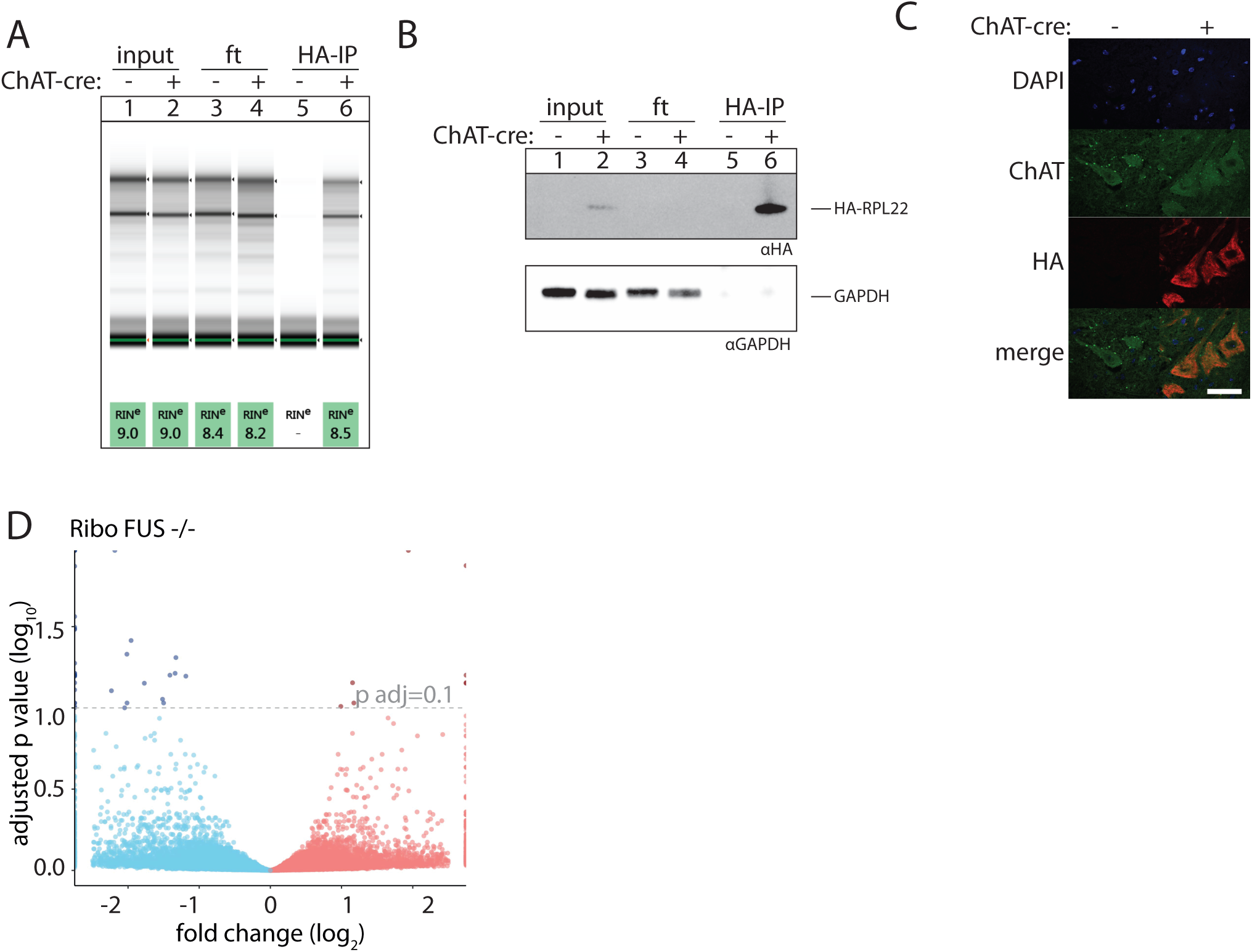
RiboTag characterisation. **(A)** TapeStation trace of RiboTag purification from adult spinal cord lysate (3 months of age) shows RNA integrity in each fraction: input (total spinal cord lysate), ft (flowthrough, unbound RNA fraction) and HA-IP (HA-bound RNA), without expression of Cre recombinase no RNA could be obtained (compare lane 5 and 6). **(B)** Western blot of fractions described in (A) show efficient precipitation of HA-RPL22. **(C)** Immunofluorescence staining of spinal cord sections of animals expressing Cre-dependent HA-RPL22 and Cre-recombinase under ChAT promoter. Compare HA labelling between Cre-and Cre+ sections. **(D)** Volcano plot of MN-specific translatome Ribo-*Fus*^-/-^. Blue points: fold change (log_2_)< 0, red points: fold change (log_2_) > 0 (n=4) (genes with fold change (log_2_) > 2.5 or <-2.5 or with p adj (-log_10_)<2 were plotted as infinity).

**SFig. 8.**
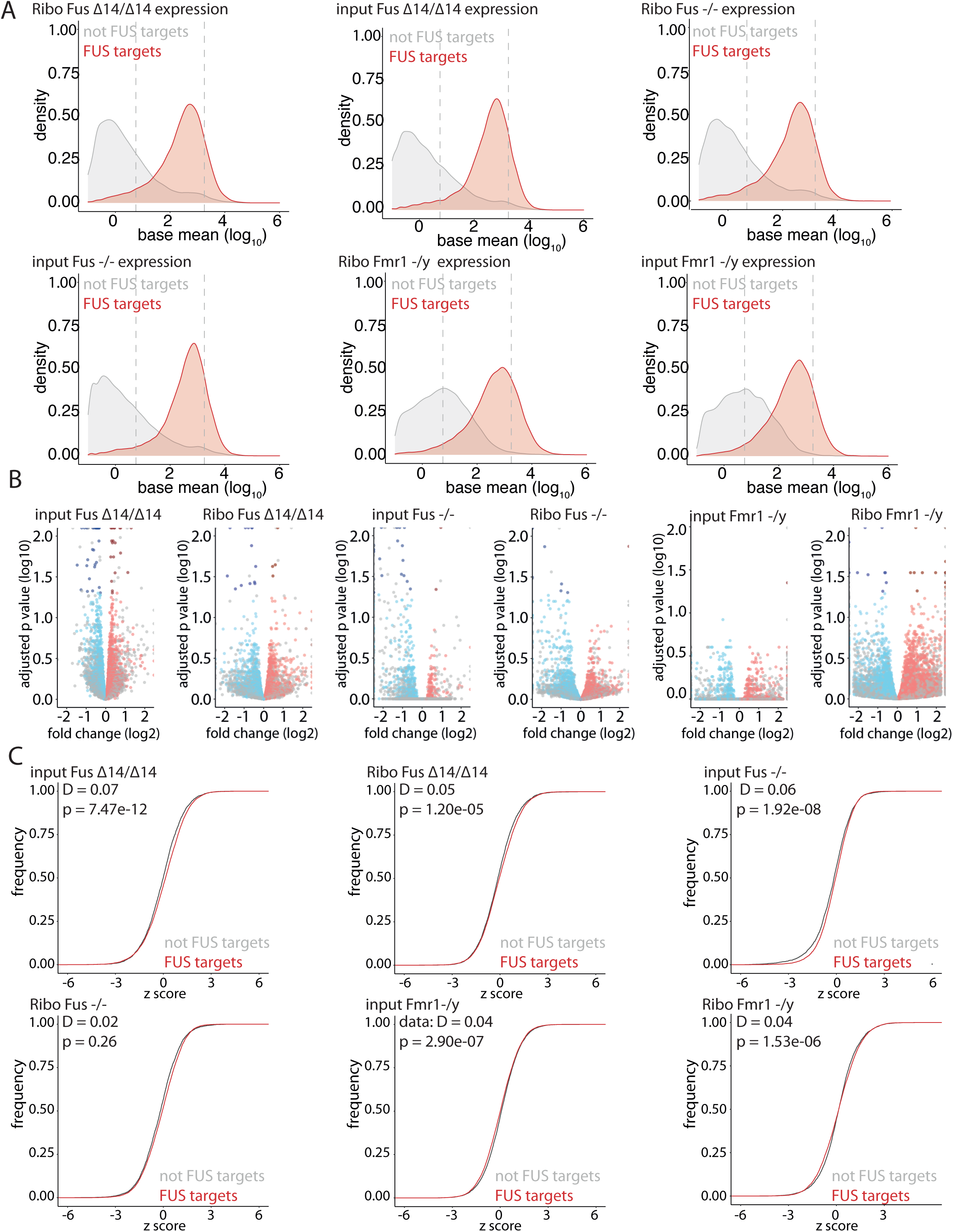
Expression of FUS targets vs transcripts without FUS binding in Ribo *Fus^Δ14/Δ14^*, *Fus^-/-^* and *Fmr1^-/y^* and their inputs. **(A)** Graphs show expression of mature FUS targets vs genes without FUS binding within the mature transcript in all datasets. Transcripts with base mean (log_10_) < 3.25 and base mean (log_10_) > 0.75 (between dashed lines) were carried forward to the next analysis. **(B)** Volcano plots of Ribosome engaged transcripts from *Fus^Δ14/Δ14^, Fus*^-/-^ and *Fmr1^-/y^* (Thomson et al., 2017), inputs from *Fus^Δ14/Δ14^, Fus*^-/-^ and *Fmr1^-/y^* (filtered by expression base mean (log_10_) < 3.25 and base mean (log_10_) > 0.75, as defined in A), FUS mature targets in red and blue; dark blue and dark red targets are significant at adjusted p value < 0.05, not FUS mature targets in grey. **(C)** Cumulative frequency plot of Z scores of transcripts shown in panel (A) (Kolmogorov-Smirnov test).

**SFig. 9.**
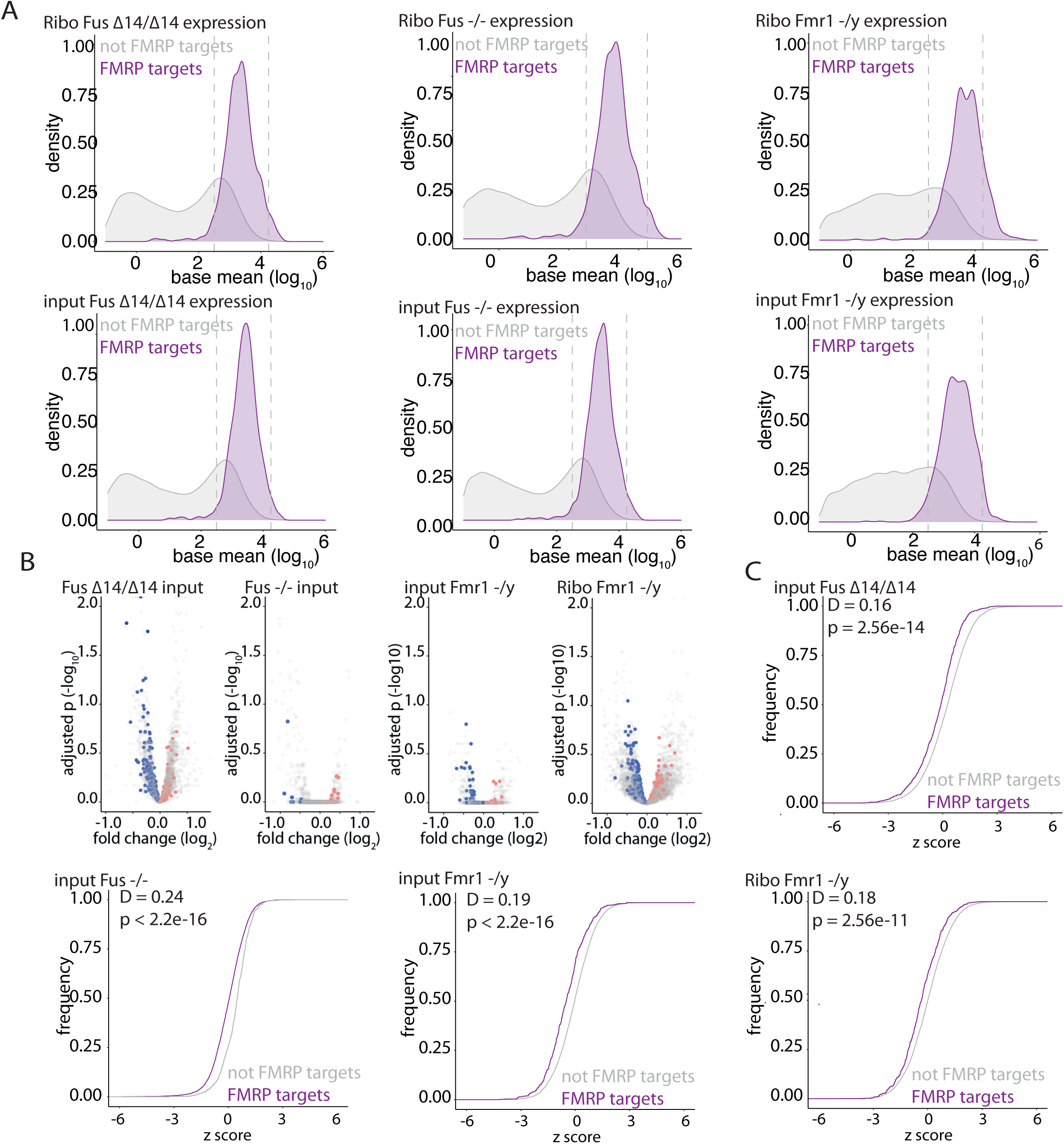
Expression of FMRP targets vs transcripts without FMRP binding in Ribo *Fus^Δ14/Δ1^*^4^, *Fus-^/-^* and *Fmr1^-/y^* and their inputs. **(A)** Graphs show expression of mature FMRP targets vs genes without FMRP binding within the mature transcript in all datasets. Transcripts with base mean (log_10_) < 4.25 and base mean (log_10_) > 2.5 (between dashed lines) were carried forward to the next analysis. Graphs show bias of expression of FMRP targets across all datasets. **(B)** Volcano plot of MN-specific translatome Ribo-*Fmr1^-/y^* (Thomson et al., 2017) and inputs from *Fus^Δ14/Δ14^, Fus*^-/-^ and *Fmr1^-/y^* filtered by expression (log_10_) base mean < 4.25 and > 2.5, FMRP targets in red and blue, not FMRP targets in grey). **(C)** Cumulative frequency plot of Z scores of genes shown in (A) show a significant decrease of FMRP targets expression: input *Fus^Δ14/Δ14^* (Kolmogorov-Smirnov test).

## Bibliography

Bernabò, P., Tebaldi, T., Groen, E.J.N., Lane, F.M., Perenthaler, E., Mattedi, F., Newbery, H.J., Zhou, H., Zuccotti, P., Potrich, V., et al. (2017). In Vivo Translatome Profiling in Spinal Muscular Atrophy Reveals a Role for SMN Protein in Ribosome Biology. Cell Rep 21, 953–965.

Birsa, N., Bentham, M.P., and Fratta, P. (2020). Cytoplasmic functions of TDP-43 and FUS and their role in ALS. Semin. Cell Dev. Biol. 99, 193–201.

Blokhuis, A.M., Koppers, M., Groen, E.J.N., van den Heuvel, D.M.A., Dini Modigliani, S., Anink, J.J., Fumoto, K., van Diggelen, F., Snelting, A., Sodaar, P., et al. (2016). Comparative interactomics analysis of different ALS-associated proteins identifies converging molecular pathways. Acta Neuropathol. 132, 175–196.

Buxbaum, A.R., Wu, B., and Singer, R.H. (2014). Single β-Actin mRNA Detection in Neurons Reveals a Mechanism for Regulating Its Translatability. Science 343, 419–422.

Carlson M, Maintainer BP (2015). TxDb.Mmusculus.UCSC.mm9.knownGene: Annotation package for TxDb object(s). R package version 3.2.2.

Carlson M (2019). org.Mm.eg.db: Genome wide annotation for Mouse. R package version 3.8.2.

Cavalcante, R.G., and Sartor, M.A. (2017). annotatr: genomic regions in context. Bioinformatics 33, 2381–2383.

Cid-Samper, F., Gelabert-Baldrich, M., Lang, B., Lorenzo-Gotor, N., Ponti, R.D., Severijnen, L.-A.W.F.M., Bolognesi, B., Gelpi, E., Hukema, R.K., Botta-Orfila, T., et al. (2018). An Integrative Study of Protein-RNA Condensates Identifies Scaffolding RNAs and Reveals Players in Fragile X-Associated Tremor/Ataxia Syndrome. Cell Rep 25, 3422–3434.e7.

Coyne, A.N., Yamada, S.B., Siddegowda, B.B., Estes, P.S., Zaepfel, B.L., Johannesmeyer, J.S., Lockwood, D.B., Pham, L.T., Hart, M.P., Cassel, J.A., et al. (2015). Fragile X protein mitigates TDP-43 toxicity by remodeling RNA granules and restoring translation. Hum. Mol. Genet. 24, 6886–6898.

Danielson, E., and Lee, S.H. (2014). SynPAnal: Software for Rapid Quantification of the Density and Intensity of Protein Puncta from Fluorescence Microscopy Images of Neurons. PLOS ONE 9, e115298.

Darnell, J.C., Van Driesche, S.J., Zhang, C., Hung, K.Y.S., Mele, A., Fraser, C.E., Stone, E.F., Chen, C., Fak, J.J., Chi, S.W., et al. (2011). FMRP stalls ribosomal translocation on mRNAs linked to synaptic function and autism. Cell 146, 247–261.

De Santis, R., Alfano, V., de Turris, V., Colantoni, A., Santini, L., Garone, M.G., Antonacci, G., Peruzzi, G., Sudria-Lopez, E., Wyler, E., et al. (2019). Mutant FUS and ELAVL4 (HuD) Aberrant Crosstalk in Amyotrophic Lateral Sclerosis. Cell Reports 27, 3818–3831.e5.

DeJesus-Hernandez, M., Kocerha, J., Finch, N., Crook, R., Baker, M., Desaro, P., Johnston, A., Rutherford, N., Wojtas, A., Kennelly, K., et al. (2010). De Novo Truncating FUS Gene Mutation as a Cause of Sporadic Amyotrophic Lateral Sclerosis. Hum Mutat 31, E1377–E1389.

Devoy, A., Kalmar, B., Stewart, M., Park, H., Burke, B., Noy, S.J., Redhead, Y., Humphrey, J., Lo, K., Jaeger, J., et al. (2017). Humanized mutant FUS drives progressive motor neuron degeneration without aggregation in “FUSDelta14” knockin mice. Brain 140, 2797–2805.

Dobin, A., Davis, C.A., Schlesinger, F., Drenkow, J., Zaleski, C., Jha, S., Batut, P., Chaisson, M., and Gingeras, T.R. (2013). STAR: ultrafast universal RNA-seq aligner. Bioinformatics 29, 15–21.

Ederle, H., and Dormann, D. (2017). TDP-43 and FUS en route from the nucleus to the cytoplasm. FEBS Lett. 591, 1489–1507.

Fallini, C., Bassell, G.J., and Rossoll, W. (2010). High-efficiency transfection of cultured primary motor neurons to study protein localization, trafficking, and function. Molecular Neurodegeneration 5, 17.

Fratta, P., Sivakumar, P., Humphrey, J., Lo, K., Ricketts, T., Oliveira, H., Brito-Armas, J.M., Kalmar, B., Ule, A., Yu, Y., et al. (2018). Mice with endogenous TDP-43 mutations exhibit gain of splicing function and characteristics of amyotrophic lateral sclerosis. EMBO J. 37.

Garone, M.G., Alfano, V., Salvatori, B., Braccia, C., Peruzzi, G., Colantoni, A., Bozzoni, I., Armirotti, A., and Rosa, A. (2020a). Proteomics analysis of FUS mutant human motoneurons reveals altered regulation of cytoskeleton and other ALS-linked proteins via 3ʹUTR binding. Scientific Reports 10, 11827.

Garone, M.G., Birsa, N., Rosito, M., Salaris, F., Mochi, M., Turris, V. de, Nair, R.R., Cunningham, T.J., Fisher, E.M.C., Fratta, P., et al. (2020b). RNA-binding protein network alteration causes aberrant axon branching and growth phenotypes in FUS ALS mutant motoneurons. BioRxiv 2020.08.26.268631.

Goering, R., Hudish, L.I., Guzman, B.B., Raj, N., Bassell, G.J., Russ, H.A., Dominguez, D., and Taliaferro, J.M. FMRP promotes RNA localization to neuronal projections through interactions between its RGG domain and G-quadruplex RNA sequences. ELife 9.

Groen, E.J.N., Fumoto, K., Blokhuis, A.M., Engelen-Lee, J., Zhou, Y., van den Heuvel, D.M.A., Koppers, M., van Diggelen, F., van Heest, J., Demmers, J.A.A., et al. (2013). ALS-associated mutations in FUS disrupt the axonal distribution and function of SMN. Hum Mol Genet 22, 3690–3704.

Hofweber, M., Hutten, S., Bourgeois, B., Spreitzer, E., Niedner-Boblenz, A., Schifferer, M., Ruepp, M.-D., Simons, M., Niessing, D., Madl, T., et al. (2018). Phase Separation of FUS Is Suppressed by Its Nuclear Import Receptor and Arginine Methylation. Cell 173, 706–719.e13.

Humphrey, J., Birsa, N., Milioto, C., McLaughlin, M., Ule, A.M., Robaldo, D., Eberle, A.B., Kräuchi, R., Bentham, M., Brown, A.-L., et al. (2020). FUS ALS-causative mutations impair FUS autoregulation and splicing factor networks through intron retention. Nucleic Acids Res.

Kamelgarn, M., Chen, J., Kuang, L., Jin, H., Kasarskis, E.J., and Zhu, H. (2018). ALS mutations of FUS suppress protein translation and disrupt the regulation of nonsense-mediated decay. Proc. Natl. Acad. Sci. U.S.A. 115, E11904–E11913.

Kim, T.H., Tsang, B., Vernon, R.M., Sonenberg, N., Kay, L.E., and Forman-Kay, J.D. (2019). Phospho-dependent phase separation of FMRP and CAPRIN1 recapitulates regulation of translation and deadenylation. Science 365, 825–829.

Kino, Y., Washizu, C., Kurosawa, M., Yamada, M., Miyazaki, H., Akagi, T., Hashikawa, T., Doi, H., Takumi, T., Hicks, G.G., et al. (2015). FUS/TLS deficiency causes behavioral and pathological abnormalities distinct from amyotrophic lateral sclerosis. Acta Neuropathol Commun 3.

Krichevsky, A.M., and Kosik, K.S. (2001). Neuronal RNA Granules: A Link between RNA Localization and Stimulation-Dependent Translation. Neuron 32, 683–696.

Lashley, T., Rohrer, J.D., Bandopadhyay, R., Fry, C., Ahmed, Z., Isaacs, A.M., Brelstaff, J.H., Borroni, B., Warren, J.D., Troakes, C., et al. (2011). A comparative clinical, pathological, biochemical and genetic study of fused in sarcoma proteinopathies. Brain 134, 2548–2564.

Lenzi, J., De Santis, R., de Turris, V., Morlando, M., Laneve, P., Calvo, A., Caliendo, V., Chiò, A., Rosa, A., and Bozzoni, I. (2015). ALS mutant FUS proteins are recruited into stress granules in induced pluripotent stem cell-derived motoneurons. Dis Model Mech 8, 755–766.

López-Erauskin, J., Tadokoro, T., Baughn, M.W., Myers, B., McAlonis-Downes, M., Chillon-Marinas, C., Asiaban, J.N., Artates, J., Bui, A.T., Vetto, A.P., et al. (2018). ALS/FTD-Linked Mutation in FUS Suppresses Intra-axonal Protein Synthesis and Drives Disease Without Nuclear Loss-of-Function of FUS. Neuron 100, 816–830.e7.

Love, M.I., Huber, W., and Anders, S. (2014). Moderated estimation of fold change and dispersion for RNA-seq data with DESeq2. Genome Biol. 15, 550.

Mejzini, R., Flynn, L.L., Pitout, I.L., Fletcher, S., Wilton, S.D., and Akkari, P.A. (2019). ALS Genetics, Mechanisms, and Therapeutics: Where Are We Now? Front. Neurosci. 13.

Mitchell, J.C., McGoldrick, P., Vance, C., Hortobagyi, T., Sreedharan, J., Rogelj, B., Tudor, E.L., Smith, B.N., Klasen, C., Miller, C.C.J., et al. (2013). Overexpression of human wild-type FUS causes progressive motor neuron degeneration in an age- and dose-dependent fashion. Acta Neuropathol. 125, 273–288.

Moens, T.G., Niccoli, T., Wilson, K.M., Atilano, M.L., Birsa, N., Gittings, L.M., Holbling, B.V., Dyson, M.C., Thoeng, A., Neeves, J., et al. (2019). C9orf72 arginine-rich dipeptide proteins interact with ribosomal proteins in vivo to induce a toxic translational arrest that is rescued by eIF1A. Acta Neuropathol 137, 487–500.

Mudge, J.M., and Harrow, J. (2015). Creating reference gene annotation for the mouse C57BL6/J genome assembly. Mamm. Genome 26, 366–378.

Murakami, T., Qamar, S., Lin, J.Q., Schierle, G.S.K., Rees, E., Miyashita, A., Costa, A.R., Dodd, R.B., Chan, F.T.S., Michel, C.H., et al. (2015). ALS/FTD Mutation-Induced Phase Transition of FUS Liquid Droplets and Reversible Hydrogels into Irreversible Hydrogels Impairs RNP Granule Function. Neuron 88, 678–690.

Neumann, M., Sampathu, D.M., Kwong, L.K., Truax, A.C., Micsenyi, M.C., Chou, T.T., Bruce, J., Schuck, T., Grossman, M., Clark, C.M., et al. (2006). Ubiquitinated TDP-43 in frontotemporal lobar degeneration and amyotrophic lateral sclerosis. Science 314, 130–133.

Neumann, M., Rademakers, R., Roeber, S., Baker, M., Kretzschmar, H.A., and Mackenzie, I.R.A. (2009). A new subtype of frontotemporal lobar degeneration with FUS pathology. Brain 132, 2922–2931.

Park, H.Y., Lim, H., Yoon, Y.J., Follenzi, A., Nwokafor, C., Lopez-Jones, M., Meng, X., and Singer, R.H. (2014). Visualization of Dynamics of Single Endogenous mRNA Labeled in Live Mouse. Science 343, 422–424.

Patel, A., Lee, H.O., Jawerth, L., Maharana, S., Jahnel, M., Hein, M.Y., Stoynov, S., Mahamid, J., Saha, S., Franzmann, T.M., et al. (2015). A Liquid-to-Solid Phase Transition of the ALS Protein FUS Accelerated by Disease Mutation. Cell 162, 1066–1077.

Phan, A.T., Kuryavyi, V., Darnell, J.C., Serganov, A., Majumdar, A., Ilin, S., Raslin, T., Polonskaia, A., Chen, C., Clain, D., et al. (2011). Structure-function studies of FMRP RGG peptide recognition of an RNA duplex-quadruplex junction. Nat. Struct. Mol. Biol. 18, 796–804.

Qamar, S., Wang, G., Randle, S.J., Ruggeri, F.S., Varela, J.A., Lin, J.Q., Phillips, E.C., Miyashita, A., Williams, D., Ströhl, F., et al. (2018). FUS Phase Separation Is Modulated by a Molecular Chaperone and Methylation of Arginine Cation-π Interactions. Cell 173, 720–734.e15.

Qiu, H., Lee, S., Shang, Y., Wang, W.-Y., Au, K.F., Kamiya, S., Barmada, S.J., Finkbeiner, S., Lui, H., Carlton, C.E., et al. (2014). ALS-associated mutation FUS-R521C causes DNA damage and RNA splicing defects. J Clin Invest 124, 981–999.

Reimand, J., Kull, M., Peterson, H., Hansen, J., and Vilo, J. (2007). g:Profiler—a web-based toolset for functional profiling of gene lists from large-scale experiments. Nucleic Acids Res 35, W193–W200.

Restani, L., Giribaldi, F., Manich, M., Bercsenyi, K., Menendez, G., Rossetto, O., Caleo, M., and Schiavo, G. (2012). Botulinum Neurotoxins A and E Undergo Retrograde Axonal Transport in Primary Motor Neurons. PLOS Pathogens 8, e1003087.

Richter, J.D., Bassell, G.J., and Klann, E. (2015). Dysregulation and restoration of translational homeostasis in fragile X syndrome. Nat. Rev. Neurosci. 16, 595–605.

Rogelj, B., Easton, L.E., Bogu, G.K., Stanton, L.W., Rot, G., Curk, T., Zupan, B., Sugimoto, Y., Modic, M., Haberman, N., et al. (2012). Widespread binding of FUS along nascent RNA regulates alternative splicing in the brain. Sci Rep 2, 603.

Sahoo, P.K., Lee, S.J., Jaiswal, P.B., Alber, S., Kar, A.N., Miller-Randolph, S., Taylor, E.E., Smith, T., Singh, B., Ho, T.S.-Y., et al. (2018). Axonal G3BP1 stress granule protein limits axonal mRNA translation and nerve regeneration. Nature Communications 9, 3358.

Sanz, E., Yang, L., Su, T., Morris, D.R., McKnight, G.S., and Amieux, P.S. (2009). Cell-type-specific isolation of ribosome-associated mRNA from complex tissues. Proc. Natl. Acad. Sci. U.S.A. 106, 13939–13944.

Scekic-Zahirovic, J., Sendscheid, O., El Oussini, H., Jambeau, M., Sun, Y., Mersmann, S., Wagner, M., Dieterlé, S., Sinniger, J., Dirrig-Grosch, S., et al. (2016). Toxic gain of function from mutant FUS protein is crucial to trigger cell autonomous motor neuron loss. EMBO J. 35, 1077–1097.

Sephton, C.F., Tang, A.A., Kulkarni, A., West, J., Brooks, M., Stubblefield, J.J., Liu, Y., Zhang, M.Q., Green, C.B., Huber, K.M., et al. (2014). Activity-dependent FUS dysregulation disrupts synaptic homeostasis. Proc. Natl. Acad. Sci. U.S.A. 111, E4769–4778.

Shigeoka, T., Jung, H., Jung, J., Turner-Bridger, B., Ohk, J., Lin, J.Q., Amieux, P.S., and Holt, C.E. (2016). Dynamic Axonal Translation in Developing and Mature Visual Circuits. Cell 166, 181–192.

Simsek, D., Tiu, G.C., Flynn, R.A., Byeon, G.W., Leppek, K., Xu, A.F., Chang, H.Y., and Barna, M. (2017). The Mammalian Ribo-interactome Reveals Ribosome Functional Diversity and Heterogeneity. Cell 169, 1051–1065.e18.

Sleigh, J.N., Tosolini, A.P., Gordon, D., Devoy, A., Fratta, P., Fisher, E.M.C., Talbot, K., and Schiavo, G. (2020). Mice Carrying ALS Mutant TDP-43, but Not Mutant FUS, Display In Vivo Defects in Axonal Transport of Signaling Endosomes. Cell Reports 30, 3655–3662.e2.

Sun, S., Ling, S.-C., Qiu, J., Albuquerque, C.P., Zhou, Y., Tokunaga, S., Li, H., Qiu, H., Bui, A., Yeo, G.W., et al. (2015). ALS-causative mutations in FUS/TLS confer gain and loss of function by altered association with SMN and U1-snRNP. Nat Commun 6, 6171.

Tan, Y., Sgobio, C., Arzberger, T., Machleid, F., Tang, Q., Findeis, E., Tost, J., Chakroun, T., Gao, P., Höllerhage, M., et al. (2020). Loss of fragile X mental retardation protein precedes Lewy pathology in Parkinson’s disease. Acta Neuropathol. 139, 319–345.

Taylor, J.P., Brown, R.H., and Cleveland, D.W. (2016). Decoding ALS: from genes to mechanism. Nature 539, 197–206.

Thomson, S.R., Seo, S.S., Barnes, S.A., Louros, S.R., Muscas, M., Dando, O., Kirby, C., Wyllie, D.J.A., Hardingham, G.E., Kind, P.C., et al. (2017). Cell-Type-Specific Translation Profiling Reveals a Novel Strategy for Treating Fragile X Syndrome. Neuron 95, 550–563.e5.

Todd, P.K., Oh, S.Y., Krans, A., He, F., Sellier, C., Frazer, M., Renoux, A.J., Chen, K., Scaglione, K.M., Basrur, V., et al. (2013). CGG repeat-associated translation mediates neurodegeneration in fragile X tremor ataxia syndrome. Neuron 78, 440–455.

Tsang, B., Arsenault, J., Vernon, R.M., Lin, H., Sonenberg, N., Wang, L.-Y., Bah, A., and Forman-Kay, J.D. (2019). Phosphoregulated FMRP phase separation models activity-dependent translation through bidirectional control of mRNA granule formation. PNAS 116, 4218–4227.

Tyzack, G.E., Luisier, R., Taha, D.M., Neeves, J., Modic, M., Mitchell, J.S., Meyer, I., Greensmith, L., Newcombe, J., Ule, J., et al. (2019). Widespread FUS mislocalization is a molecular hallmark of amyotrophic lateral sclerosis. Brain 142, 2572–2580.

Vance, C., Rogelj, B., Hortobágyi, T., Vos, K.J.D., Nishimura, A.L., Sreedharan, J., Hu, X., Smith, B., Ruddy, D., Wright, P., et al. (2009). Mutations in FUS, an RNA Processing Protein, Cause Familial Amyotrophic Lateral Sclerosis Type 6. Science 323, 1208–1211.

Wang, J., Choi, J.-M., Holehouse, A.S., Lee, H.O., Zhang, X., Jahnel, M., Maharana, S., Lemaitre, R., Pozniakovsky, A., Drechsel, D., et al. (2018). A Molecular Grammar Governing the Driving Forces for Phase Separation of Prion-like RNA Binding Proteins. Cell 174, 688–699.e16.

Yamazaki, T., Chen, S., Yu, Y., Yan, B., Haertlein, T.C., Carrasco, M.A., Tapia, J.C., Zhai, B., Das, R., Lalancette-Hebert, M., et al. (2012). FUS-SMN Protein Interactions Link the Motor Neuron Diseases ALS and SMA. Cell Reports 2, 799–806.

Zhou, Y., Liu, S., Liu, G., Öztürk, A., and Hicks, G.G. (2013). ALS-Associated FUS Mutations Result in Compromised FUS Alternative Splicing and Autoregulation. PLOS Genetics 9, e1003895.

